# Permanent Neonatal diabetes-causing Insulin mutations have dominant negative effects on beta cell identity

**DOI:** 10.1101/2023.09.01.555839

**Authors:** Yuwei Zhang, Lina Sui, Qian Du, Leena Haataja, Yishu Yin, Ryan Viola, Shuangyi Xu, Christian Ulrik Nielsson, Rudolph L. Leibel, Fabrizio Barbetti, Peter Arvan, Dieter Egli

## Abstract

Heterozygous coding sequence mutations of the *INS* gene are a cause of permanent neonatal diabetes (PNDM) that results from beta cell failure. We explored the causes of beta cell failure in two PNDM patients with two distinct *INS* mutations. Using b and mutated hESCs, we detected accumulation of misfolded proinsulin and impaired proinsulin processing *in vitro*, and a dominant-negative effect of these mutations on the in vivo performance of patient-derived SC-beta cells after transplantation into NSG mice. These insulin mutations derange endoplasmic reticulum (ER) homeostasis, and result in the loss of beta-cell mass and function. In addition to anticipated apoptosis, we found evidence of beta-cell dedifferentiation, characterized by an increase of cells expressing both Nkx6.1 and ALDH1A3, but negative for insulin and glucagon. These results highlight both known and novel mechanisms contributing to the loss and functional failure of human beta cells with specific insulin gene mutations.

## Introduction

Neonatal Diabetes Mellitus (NDM) is a rare disorder with an incidence of about 1/100,000 live births; approximately 50% of cases are permanent (PNDM) (Iafusco et al., 2012). Heterozygous coding sequence mutations of the insulin (*INS*) gene are among the leading causes of PNDM (Colombo et al., 2008; De Franco et al., 2015; Edghill et al., 2008; Støy et al., 2007). In Western Europe and the U.S., mutations in *KCNJ11*, *INS* and *ABCC8* account for about 65-70% of PNDM cases (Greeley et al., 2022). Other causes are structural abnormalities on chromosome 6q24, as well as biallelic *GCK* and *PDX1* mutations, but the latter are exceedingly rare (De Franco et al., 2015). In the Middle East, the leading gene is *EIF2AK3* causing Wolcott-Rallison syndrome, followed by mutations in other recessive genes, including *PTF1A* and biallelic recessive *INS* mutations (Habeb et al., 2012). PNDM is usually diagnosed within 6 months of birth, presenting variously as mild hyperglycemia with failure to thrive or fatal diabetic ketoacidosis (DKA) (Colombo et al., 2008; Dahl and Kumar, 2020; Edghill et al., 2006; Iafusco et al., 2002; Redondo et al., 2020).

Diabetic mouse models segregating for dominant *INS* gene mutations, such as the Akita (*INS2*^+/C96Y^) and Munich (*INS2*^+/C95S^) mice have been studied to elucidate the molecular pathogenesis of the resulting diabetes, as their early-onset diabetic phenotypes resemble human PNDM (Herbach et al., 2007; Izumi et al., 2003; Oyadomari et al., 2002; Wang et al., 1999; Yoshioka et al., 1997). Disrupted proinsulin folding and activated endoplasmic reticulum (ER) stress have been implicated in these heterozygous mutant mice. Transfection of mutated insulin transcripts nominally affecting protein folding in beta cell lines causes cell death (Colombo et al., 2008). In all of these models, the presence of a non-mutant *INS* allele does not rescue the beta cell failure phenotype (Wright et al., 2013).

Proinsulin folding occurs primarily within the ER. Native proinsulin folding is associated with the formation of three specific disulfide bonds pairing 6 cysteines (A6-A11, A7-B7, and A20-B19; A: A chain, B: B-chain) (Weiss, 2009). Conserved among most members of the insulin superfamily, these disulfide bridges are critical to the 3-dimensional structure of proinsulin and the bioactivity of insulin (Liu et al., 2018). *INS* mutations affecting cysteine residues can directly interfere with proper proinsulin disulfide bond formation (Colombo et al., 2008; Ichi Nozaki et al., 2004; Yoshinaga et al., 2005); and various non-cysteine mutations can also impair cysteine alignment leading to defective disulfide bond formation (Colombo et al., 2008; Haataja et al., 2021; Herbach et al., 2007). Increased accumulation of misfolded proinsulin invokes ER chaperone proteins such as BIP/GRP78 and GRP94 (Liu et al., 2018; Oyadomari et al., 2002). Pancreatic beta-cells are susceptible to perturbations of ER homeostasis which can cause cell death in mice with insulin mutations (Oyadomari et al., 2002), T2D and possibly T1D (Brozzi and Eizirik, 2016). However, direct proof that ER stress triggers programmed beta-cell death in diabetic patients (or in PNDM infants) is lacking (Ron, 2002; Yong et al., 2021).

Patient-derived human induced pluripotent stem cells (hiPSCs) can be differentiated to pancreatic beta-cells (SC beta-cells), offering the prospect of replacement cell therapy for both T1D as well as for monogenic forms of diabetes (Brusko et al., 2021; Johannesson et al., 2015). Many studies have shown the value of stem cell models of monogenic forms of diabetes for gaining understanding of the pathophysiology of more prevalent types of diabetes mellitus (Abdelalim, 2020).

Endoplasmic reticulum (ER) stress affects insulin biosynthesis and secretion in stem cell derived beta cells (Balboa et al., 2018; Maxwell et al., 2020; Shang et al., 2014). In the present study, we sought to understand mechanisms driving beta cell failure associated with two distinct pathologic *INS* mutations: L^B15^Y^B16^delinsH and Y^B26^C (current nomenclature according to Human Genome Variation Society: p.Leu39_Tyr40delinsHis and p.Tyr50Cys), using differentiation of human iPSCs and embryonic stem cells (hESCs). We corrected patient-derived iPSCs using CRISPR/Cas9 to create isogenic controls, along with WT iPSCs and hESCs as additional controls. Differentiation efficiency, and short-term proinsulin/insulin synthesis and proinsulin foldability were examined *in vitro*. Development and maintenance of beta-cell identity, as well as ER stress, were assessed *in vivo* upon transplantation of stem cell-derived beta cells into immunodeficient mice. Collectively, our findings indicate that specific heterozygous *INS* mutations in human beta cells cause diabetes by dominant-negative effects on insulin biosynthesis and secretion, as well as through a newly identified mechanism: de-differentiation and loss of beta-cell identity.

## Results

### 2 patients segregating for *INS* mutations with distinct neonatal diabetes syndrome

A female neonate (Policlinico Tor Vergata **PTV1**) born at term (birth weight: 3350 g) after an uneventful pregnancy was diagnosed with diabetic ketoacidosis (DKA) at 37 days of age; plasma glucose 613 mg/dl and arterial blood pH 7.22 (Colombo et al., 2008, kindred D) (**Figure 1A**). T1DM-related autoantibody status is unknown. Plasma C-peptide was low but measurable up to one year of age and subsequently became undetectable. Insulin administration was initiated at the time of diagnosis (d37) with a full dose at 1 UI/kg/d with good metabolic control (mean HbA1c 7.3% from 1994-2013). Currently at age 24, she remains free of diabetic complications (**Figure 1A**) (Iafusco et al., 2014, case #2).

**Figure 1.**
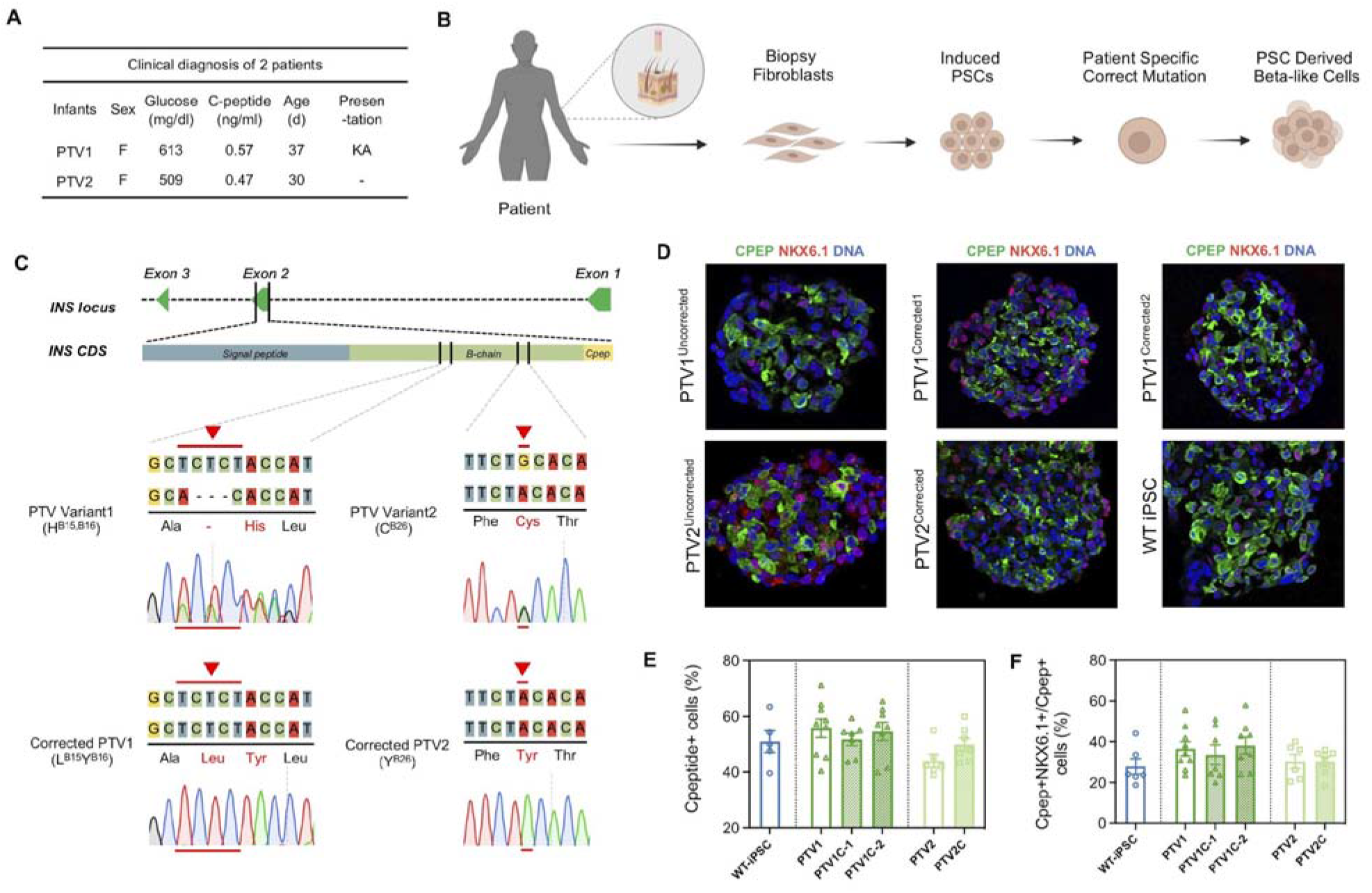
Generation of neonatal diabetes disease models with insulin mutations and their isogenic controls. **(A)**. Diagnosis of two permanent neonatal diabetes (PNDM) patients segregating formutations in the insulin gene (*INS*). Age at diagnosis is shown, with corresponding glucose levels, and symptoms. C-peptide concentrations were measured at day 40 for PTV1 and day 37 for PTV2. PTV: Policlinico Tor Vergata patient variant; F: female; KA: Ketoacidosis. **(B)**. A schematic of the PNDM model. Patient fibroblasts are reprogrammed to iPS cells, isogenic controls were made by gene correction, and both mutant and corrected cells were differentiated to beta-like cells. **(C)**. Molecular identity of patient variants PTV1 and PTV2 and CRSPR/Cas9-corrected isogenic controls were examined by DNA sequencing. **(D)**. Immunostaining of frozen sections of PTV1 and corrected PTV1C, PTV2 and PTV2C, and WT iPSC-derived beta cell clusters at d27 for C-peptide (the antibody detects both C-peptide and aa 33-63 of proinsulin), NKX6.1. scale bar: 50μm. **(E)**. Quantitation of beta cell differentiation efficiency indicated by C-peptide+ population (%) and **(F)** C-peptide+ cells co-expressing NKX6.1 population (%). n= 5-9 independent differentiation experiments per genotype. Respective cell numbers quantified in cell blocks with a mean of ∼130 cells per block) and presented as mean ± SEM. One-way ANOVA with *: P<0.05;

A second patient (**PTV2**; a female with a birth weight of 2500 g) was diagnosed with diabetes (plasma glucose 509 mg/dl) at one month of age, without accompanying ketoacidosis or other complications (Ortolani et al., 2016); plasma C-peptide (0.47 ng/l). The patient was treated with continuous subcutaneous insulin infusion (0.7 UI/kg/d) and discharged uneventfully. Good metabolic control without complications (mean HbA1c 6.4%) has been maintained up to the present age of 8 years (**Figure 1A** and **Table S1**). However, C-peptide levels were halved within six months of diagnosis, and were barely detectable by age 6 years. The phenotype of both patients is consistent with declining function of beta cells over time.

### *INS* mutant iPSC lines efficiently generate insulin-producing cells

We generated patient-specific iPSC cell lines by reprogramming skin fibroblasts of the two patients (designated PTV1 and PTV2; **Figure 1B**). The patients carried distinct heterozygous mutations: an in-frame substitution in which B-chain Leu15 and Tyr16 are replaced by a single His residue (L^B15^Y^B16^delinsH; **PTV1**); a point mutation in which B-chain Tyr26 is replaced by Cys (Y^B26^C, adding an additional cysteine to the B chain **PTV2**). We established 3 isogenic control cell lines by correcting the mutation of patient-derived iPSCs with CRISPR/Cas9: 2 corrected cell lines edited from **PTV1** (denominated **PTV1C-1** and **PTV1C-2**), and 1 corrected cell line from PTV2 (denominated **PTV2C**) (**Figure 1C**).

We differentiated 6 iPSC lines: 2 patient-derived iPSCs, 1 non-isogenic healthy control WT-iPSCs (line # 1159, from donors without diabetes) and 3 isogenic control iPSC lines into SC-derived beta cells using a previously published protocol (Sui et al., 2018b). The differentiation protocol is outlined in **Figure S1A**. All cell lines were differentiated into islet-like clusters containing insulin-expressing cells and were karyotypically normal (**Figure S1B**, **C**).

We characterized the islet-like clusters by immunochemistry and flow cytometry for beta cell markers: C-peptide and NKX6.1. Equivalent percentages (∼50%) of C-peptide-positive cells were detected among all the cell lines (**Figure 1D, E** and **Figure S1B**). For **PTV1**, 36% of cells co-expressed both C-peptide+ and NKX6.1, similar to isogenic controls including **PTV1C-1** (34%) and **PTV1C-2** (38%). The percentage of NKX6.1+/C-peptide+ double-positive cells in **PTV2** patient cell lines with or without correction was 30% and not significantly different from other lines (**Figure 1D, F** and **Figure S1B**). These data suggest that PTV1 and PTV2 mutations do not affect cellular developmental ability to generate pancreatic beta-like cells.

### Defective proinsulin processing, insulin formation, and secretion in patient-derived beta cells

To determine if these *INS* mutations affect insulin production in differentiated stem cell derived beta cell clusters on d27, we examined proinsulin and insulin levels by immunoassay in PTV1 and PTV2 cells, their isogenic controls (PTV1C-1, PTV1C-2, and PTV2C) and WT iPSCs. PTV1C-1, PTV1C-2 cells produced more proinsulin and insulin compared to PTV1 and PTV2 cells (**Figure 2A**, **B**). We also evaluated expression of proinsulin and insulin in SC-clusters by immunostaining (**Figure S2A**). A similar percentage of proinsulin positive cells was detected in each genotype (**Figure S2B**). However, the intensity of insulin expression was reduced in SC-clusters with PTV1 mutation (H^B15, B16^) compared to isogenic controls, and a similar trend was observed in clusters with the PTV2 mutation (C^B26^) (**Figure S2C**). The ratio of proinsulin to insulin immunopositive cells was significantly higher in both patient-derived sc-clusters, suggesting that these mutations may be associated with deficient insulin formation from proinsulin (**Figure S2D**).

**Figure 2.**
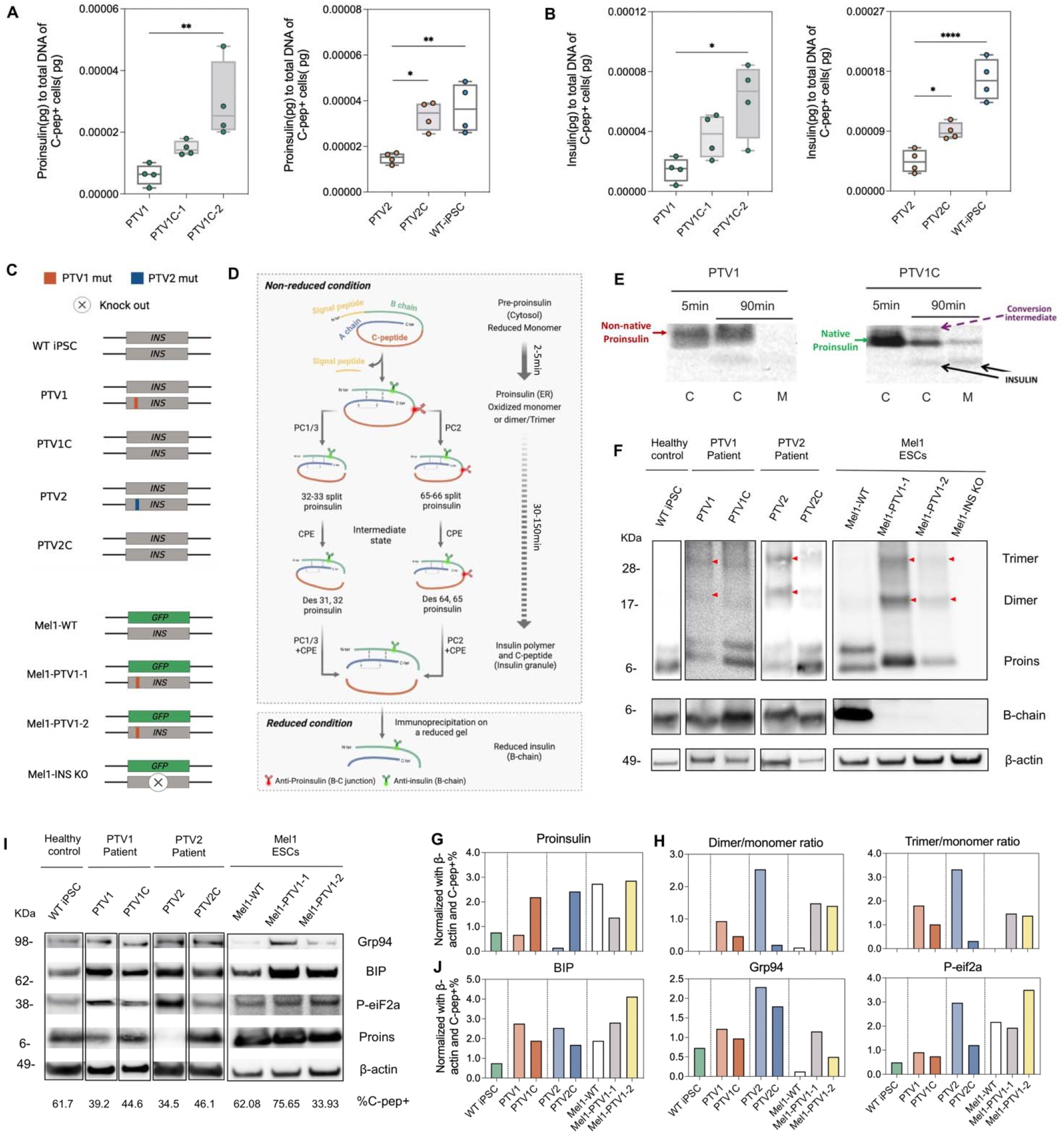
Reduced Proinsulin and insulin production in *INS* mutants accompanied by elevated ER stress and UPR. **(A)** Human proinsulin content and **(B)** insulin content in sorted SC-beta cells clusters derived from WT iPSC, PTV mutants (PTV1 and PTV2) and their isogenic controls (PTV1C-1, PTV1C-2 and PTV2C). Content was normalized to the percentage of insulin-positive cells determined by flow cytometry. n=4 per genotype. Data plots are presented as mean ± SEM. One-way ANOVA test with *P<0.05, **P < 0.01, ***P < 0.001, ****P < 0.0001. **(C)** Genotypes of iPSC lines and Mel1-hESC lines. Insulin-expressing allele is shown in grey; GFP allele is shown in green. PTV1 and PTV2 mutations are indicated respectively by orange and blue bars. **(D)** Schematic of insulin processing. Uncleaved pre-proinsulin in a reduced monomer form is processed to proinsulin, which is then cleaved to intermediates in the ER visible as double proinsulin bands on a non-reducing gel in the process of cleavage of proinsulin to insulin. Intermediate state: The junctions of the B and A chains, between residues 32 and 33 (R↓E) and residues 65 and 66 (R↓G) are recognized by the neuroendocrine convertases PC1/3 and PC2, respectively, during initial processing cleavages to form split-proinsulin. The resulting cleavage products extend insulin at the B chain C terminus through R31-R32 and extend C-peptide C-terminally through K64-R65. Subsequently, carboxypeptidase E (CPE) trims these products by removing the basic residues to form des-proinsulin (Top). A reducing gel shows only the processed B chain with the antibody used (Bottom). Illustrated using Biorender. **(E)** Pulse-chase experiment using non-reducing gel. Western blot for insulin using an antibody specific to B-chain for cellular content and secretion. Misfolded non-native proinsulin with abnormal disulfide linkages (red arrow), normally folded proinsulin (green arrow, below), conversion intermediate of proinsulin to insulin (purple arrow) and insulin (black arrow). C: samples collected from cells; M: samples collected from media to detect secreted insulin. Note that only gene-corrected cells process and secrete insulin efficiently. **(F)** Western blots showing the processing of non-reduced proinsulin (top) and distinct higher molecular weight bands corresponding to dimeric and trimeric proinsulin (dimer and trimer) indicating intermolecular disulfide bond formation in mutants. Anti-proinsulin antibody specifically detecting B-C junction sequence KTRREAEDLQ was used to detect non-reduced proinsulin. This antibody detects a mutant protein in MEL-1cells.The expression of processed B-chain recognized by an anti-Insulin antibody binding on B-chain is shown using a reducing gel below. **(G)** Quantitation of Western blot data. Proinsulin is normalized with β-actin and differentiation efficiency quantified by flow cytometry for the percentage of C-peptide positive cells (%C-pep+). **(H)** Trimer/monomer and dimer/monomer ratio as an indication of proinsulin to insulin processing failure in PTV mutant cells. **(I)** Representative Western blots showing ER stress markers including Grp94, BIP, P-eiF2a and proinsulin expression in all lines. Normalized with β-actin. Lanes are denominated by their origins (WT iPSCs from a healthy control. PTV1 and its isogenic control PTV1C from patient carrying PTV1 mutation, PTV2 and PTV2C from another patient with PTV2 mutation. WT-MEL1, Mel1-PTV1-1 and Mel1-PTV1-2 from Mel1-hESC). **(J)** Quantitation of Western blot for ER stress proteins.

We introduced the PTV1 mutation (H^B15, B16^) into the Mel1 human embryonic stem cell line (**Mel1-WT**) and generated two hemizygous cell lines (**Mel1-PTV1-1** and **Mel1-PTV1-2**) and one insulin knock out cell line as a negative control (**Mel1-INS KO**). Several attempts to introduce the PTV2 mutation into the Mel1 ESC line were unsuccessful. Genotypes of the 4 cell lines used are illustrated in **Figure 2C**, and **Figure S3A**. Because Mel1 cells carry an inactivating GFP knock-in on one of the insulin alleles, Mel1-derived cell lines express only a mutant protein, while patient cells express both a wild type and a mutant protein (**Figure 2C, Figure S3A-E**). These mutant Mel1 cell lines differentiated to beta-like cells with efficiency comparable to that of unmanipulated Mel1 cells, as quantified by flow cytometry using an antibody recognizing C-peptide (including cleaved C-peptide and aa 33-63 of proinsulin) and Nkx6.1, and GFP fluorescence (**Figure S3F, G**). Insulin knockout cell lines expressed GFP, but no insulin (**Figure S3H**). All cell lines had normal karyotypes (**Figure S3I**). Similar to patient iPSCs, Mel1 cells carrying a PTV1 mutation were immunopositive for proinsulin, and only weakly insulin immunopositive (**Figure S3J, K**).

To assess proinsulin processing in the mutant cells we performed Western blotting for processing intermediates. Normally, upon excision of the preproinsulin signal peptide in the endoplasmic reticulum, proinsulin disulfide bridges form between specific cysteine residues, followed by intracellular transport of proinsulin to secretory granules in which proinsulin is cleaved by PCSKs to generate mature insulin (Liu et al., 2014) (**Fig. 2D**).

We quantified synthesis of proinsulin, its processing to insulin and the secretion of insulin. Stem cell derived beta cells (sc-beta) PTV1 cells were pulse-labeled using ^35^S-amino acids, and chased for either 5 min or 90 min. At 5 and 90 min, proinsulin and insulin were immunoprecipitated with anti-insulin and analyzed by nonreducing SDS-PAGE. Whereas gene-corrected PTV1C cells formed native proinsulin, the uncorrected patient cells exhibited a proinsulin band of abnormal mobility (i.e., non-native), detected after both 5 min and 90 min of chase (**Figure 2E**). Additionally, whereas the gene-corrected cells synthesized and secreted insulin, in the PTV1 patient cells, formation and secretion of labeled insulin was markedly impaired (**Figure 2E**). These data are consistent with defects in proinsulin folding and insulin formation in PTV1 sc-derived beta cells.

Misfolding of proinsulin monomers can promote the formation of aberrant intermolecular proinsulin disulfide bonds (Alam et al., 2021; Arunagiri et al., 2019; Haataja et al., 2021; Sun et al., 2020). We examined this possibility using anti-proinsulin immunoblotting of protein extracts from stem cell derived beta cell clusters, and resolved by nonreducing SDS-PAGE. All genotypes were examined, including normal controls, heterozygous mutant, and corrected patient cells for both variants, as well as PTV1 hemizygous mutant ESCs and controls. In cells carrying at least one copy of WT *INS* (WT iPSC, PTV1, PTV1C, PTV2, PTV2C and Mel1-WT), proinsulin was detected in two bands near the 6 kD molecular mass marker, presumably representing intact proinsulin (lower band) and a proinsulin conversion intermediate (upper band), a pattern consistent with proinsulin folding, trafficking, and ongoing maturation in the secretory pathway (**Figure 2F**). In contrast, no normal proinsulin (or intermediate) bands were found in samples carrying only the mutant *INS* copy. These had only a proinsulin band of slightly abnormal mobility in Mel1-PTV1-1 and Mel1-PTV1-2), consistent with defective proinsulin monomer folding and an inability of that proinsulin to undergo normal trafficking and maturation in the secretory pathway (**Figure 2F**). Consistent with these findings, by reducing SDS-PAGE, cells containing a wild type copy showed the presence of a processed insulin B-chain, confirming the ability of proinsulin to undergo trafficking and maturation in the secretory pathway. Whereas in cells bearing only a mutant PTV1 allele, this pattern was not observed (**Figure 2F**). Thus, the variant PTV1 mutation (H^B15, B16^) is incompatible with the formation of mature insulin.

We also detected aberrant disulfide-linked proinsulin dimers and trimers within patient-derived and Mel1-derived beta-like cells bearing the PTV1 *INS* mutation (**Figure 2F**). Though these complexes were also seen in wild type cells, PTV1 cells showed decreased proinsulin monomer per insulin-positive cell (**Figure 2G**), and an increased ratio of proinsulin trimer and dimer to monomer, especially in the sc-derived beta-like cells expressing the proinsulin mutant (PTV2) bearing the extra cysteine residue **(Figure 2H**). These aberrant signals are allele-specific, as insulin knockout cells do not show either monomeric or oligomeric proinsulin signals despite their ability to differentiate (**Figure 2F**).

These data demonstrate that the PTV mutations interfere with proper formation of disulfide bonds and increase aberrant disulifide-linked proinsulin dimers and trimers, resulting in diminished proinsulin endoproteolytic processing due to defective anterograde proinsulin trafficking in the secretory pathway. These derangements contribute to reduced insulin production in *INS* mutant cells.

### Increased ER stress in insulin mutant sc-beta cells

Retention of proinsulin in the ER has previously been reported in Akita mice segregating for a tertiary structure-altering missense mutation (Cys96Tyr) of *INS*2 (Colombo et al., 2008; Oyadomari et al., 2002). We assessed ER stress in sc-derived beta cell clusters (at 27d of in vitro differentiation) by assaying protein levels of BIP, GRP94 and phospho-eiF2α. By Western blotting, these ER stress markers were upregulated in insulin-mutant cells with PTV1 and PTV2 mutations (**Figure 2I**, **J)**. Specifically, in patient iPSC-derived beta cell clusters carrying PTV1 and PTV2 mutations, immunofluorescence of BIP was increased in insulin-positive cells (**Figure S4A, C**). Likewise, Mel1 cells carrying a PTV1 mutation also showed an increased number of BIP/insulin double-positive cells (**Figure S4B, D**). These data suggest that - as in Akita mouse beta cells - impaired folding of proinsulin results in increased ER stress which decreases insulin production (Lipson et al., 2006).

### Reduced C-peptide secretion in patient-derived beta cell grafts transplanted in NSG mice

We tested the function of patient-derived SC-beta cells *in vivo*, by grafting these cells into immunodeficient NSG mice. We transplanted 1∼2×10^6^ beta cells from mutant, isogenic gene-corrected controls, and WT controls, into the gastrocnemius muscle of NSG mice. Beta cells grafted into mice display physiologic glucose-stimulated insulin secretion, allowing detailed functional interrogation of specific beta cell genes (Sui et al., 2021). Here, we monitored human C-peptide secretion in the fed state and after intraperitoneal glucose challenge.

In mice engrafted with WT iPSC (1159)-derived or ESC (Mel1)-derived cells, interim serum concentrations of human C-peptide in the random (fed) state rose gradually to 1248 pM in Mel1-WT engrafted mice or 1210 pM in WT-iPSC (**Figure 3A**). This time course suggests that the SC-derived beta cells integrated with the mouse vascular system and were maturing *in vivo*. In contrast, NGS mice transplanted with PTV1 or PTV2 grafts showed detectable but extremely low random C-peptide concentrations post-transplantation (**Figure 3A**). We also examined random mouse C-peptide concentrations at 7 months after grafting. As expected, secretion of mouse C-peptide in WT SC-islets engrafted mice was suppressed due to insulin secretion from the human beta cell grafts. In PTV SC-islets engrafted mice, high levels of mouse C-peptide persisted, consistent with a failure of human insulin secretion in these animals (**Figure 3B, C)**. Thus, whereas a single wild type insulin gene is sufficient in MEL-1 cells to regulate mouse blood glucose levels, cells with a single wild type insulin allele in combination with a mutant insulin allele fail to secrete physiologically sufficient amounts of insulin. These data are consistent with a dominant-negative impact of structural mutations of proinsulin molecules on the function of patient-derived SC-beta cells *in vivo*.

**Figure 3.**
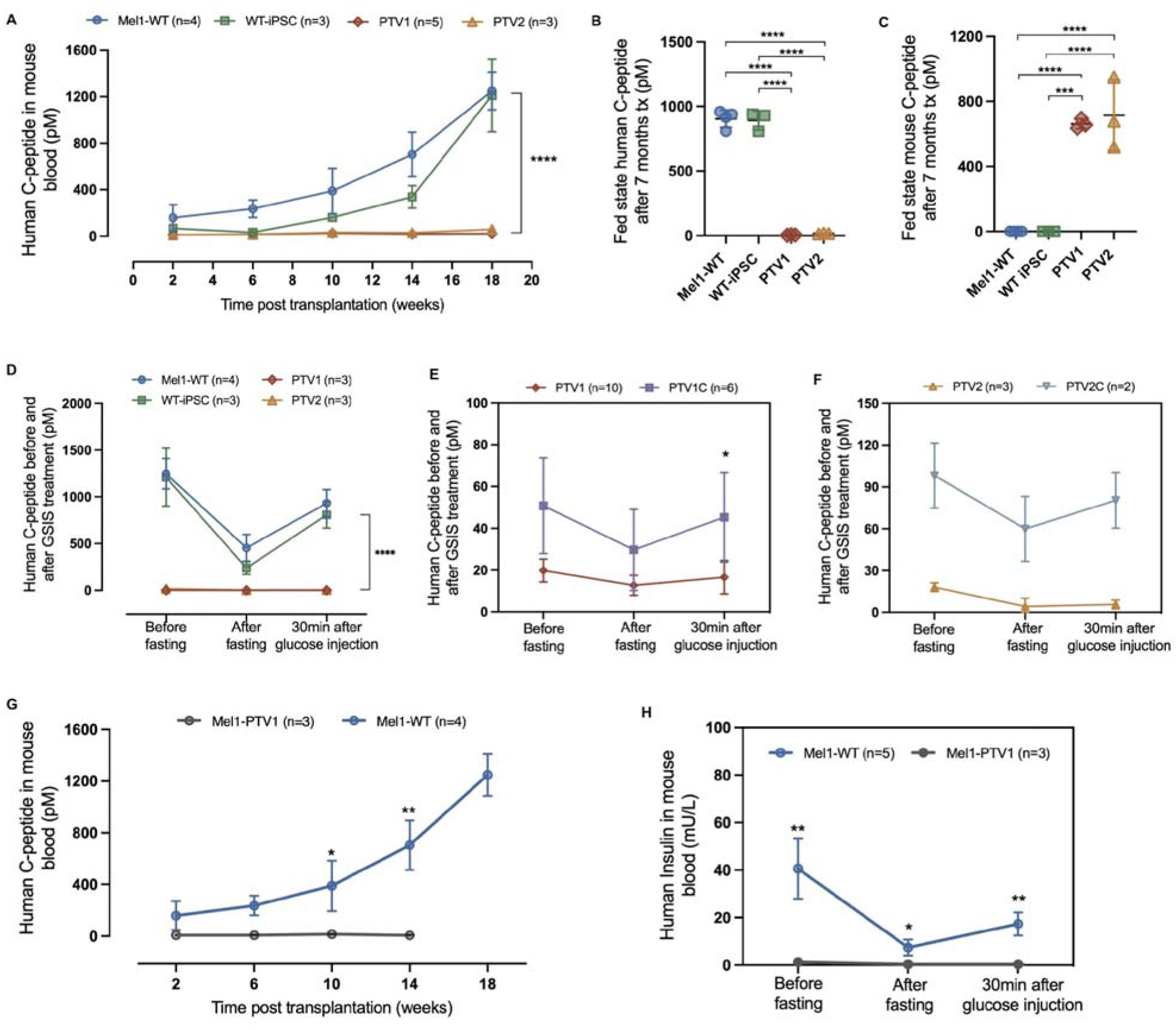
PTV SC-islets fail to secrete insulin and are unable to to regulate blood glucose levels in mice. **(A)** Human specific C-peptide levels(pM) in WT-iPSC (INSwt/wt), Mel1-WT (INSgfp/wt), PTV1 (INSptv1/wt) and PTV2 (INSptv2/wt) in plasma of mice engrafted with SC-islets. Human **(B)** and Mouse **(C)** fed state C-peptide levels(pM) after 7 months of transplantation. (Tx: Transplantation). Individual data points presented. One-way ANOVA and Tukey’s multiple comparisons test. **(D-F)** Human C-peptide (pM) measured during glucose-stimulated insulin secretion (GSIS) at 7 months post-transplantation. Mice were injected IP with 2g of glucose/kg body mass after overnight fasting (16 hours). **(G)** Measurements of human specific C-peptide levels(pM) in Mel1-WT (INS^+/GFP^) and Mel1-PTV1 (INS^PTV1/GFP^) SC-islets engrafted mice blood after transplantation. **H)** Human Insulin (mU/L) measured during glucose-stimulated insulin secretion (GSIS) at 7 months post-transplantation. Mice were injected IP with 2g of glucose/kg body mass after overnight fasting (16 hours). Data plots are presented as mean ± SEM. Two-way ANOVA test with *P<0.05, **P < 0.01, ***P < 0.001, ****P < 0.0001.

To further assess the metabolic physiology of mice transplanted with beta cells expressing mutant or corrected *INS*, mice were fasted overnight and then challenged with intraperitoneal glucose (2g/kg body mass). Mice engrafted with PTV1 or PTV2 cells displayed impaired human C-peptide secretion after glucose stimulation, compared to mice engrafted with wild type ESC and iPSC controls (**Figure 3D**). Importantly, mice with PTV1C and PTV2C gene-corrected transplants showed physiological decreases of circulating human C-peptide during fasting and increased C-peptide in response to glucose, while mice with uncorrected mutant cells did not (**Figure 3E, F**). However, the absolute serum concentration of human C-peptide in mice with gene-corrected cells did not reach those of MEL-1 cells, and were not sufficient to fully substitute for the endogenous mouse beta cells (**Figure S5A**). We attribute this partial substitution to the variability in iPSC quality (Sui et al., 2018b). Importantly, we did not ablate mouse beta cells in any of these studies. Streptozotocin can be dosed to preferentially kill endogenous mouse beta cells, but not transplanted human cells (Estilles et al., 2018; Tuch et al., 1989) Here, we were concerned that the proposed increased ER stress-susceptibility of the mutant cells would render them more sensitive to the STZ, confounding subsequent inferences regarding effects of the mutations on cell-autonomous in vivo functions of these cells.

We also measured serum glucose concentrations during intraperitoneal GSIS. Mice transplanted with PTV cells had higher serum glucose concentrations at 30 min post injection than mice with either MEL-1 grafts or with control iPS grafts (**Figure S5B**). Random glucose concentrations were lower in mice engrafted with control grafts derived from Mel1-WT/WT-iPSC compared to those bearing PTV grafts (**Figure S5C**).

We also transplanted NSG mice with Mel1 hESCs carrying the PTV1 mutation; controls were grafted with Mel1-WT cells (**Figure 2C**). As noted, Mel1 cells have only one functional *INS* allele as the other allele is null by virtue of insertion of GFP. Hence the Mel1 cells are all hemizygous for *INS*. No circulating human C-peptide was detected in mice engrafted with Mel1-PTV1 cells (**Figure 3G**), and no human insulin was detected during an GSIS during which wild type grafts produced concentrations of insulin (**Figure 3H**). These data indicate that the PTV1 allele does not produce secretable insulin (**Figure 3C**).

### Loss of *INS* mutant beta-like cells after transplantation is associated with ER stress

We assessed the impact of nominal ER stress on survival of PTV1 and PTV2 beta cells after grafting. We compared grafted PTV1 and PTV2 beta cells with their corresponding corrected and wild-type control beta cells at 7 months post transplantation. We observed a severe reduction in the percentage of C-peptide-expressing cells in the mutants, as well as the intensity of immunostaining for insulin (**Figure 4A, B** and **Figure S6A, B**). However, at the time of initial transplantation, these alterations had not yet occurred, as the differentiation efficiency remained equivalent between mutant subjects and control groups (**Fig. 1E, F**). Interestingly, most of the limited number of C-peptide-expressing cells in PTV1 and PTV2 grafts were immunopositive for the ER stress marker BIP, in contrast to much lower frequency in gene-corrected beta cells or uncorrected control cells (**Figure 4A, C** and **Figure S6A, C**). Specifically, 64% and 67%, respectively of PTV1 and PTV2 C-peptide positive cells co-expressed BIP. Such double-positive cells were much rarer in control (3.50%) and corrected isogenic PTV1C (8.28%) and PTV2C (0.36%) cells (**Figure 4C**).

**Figure 4.**
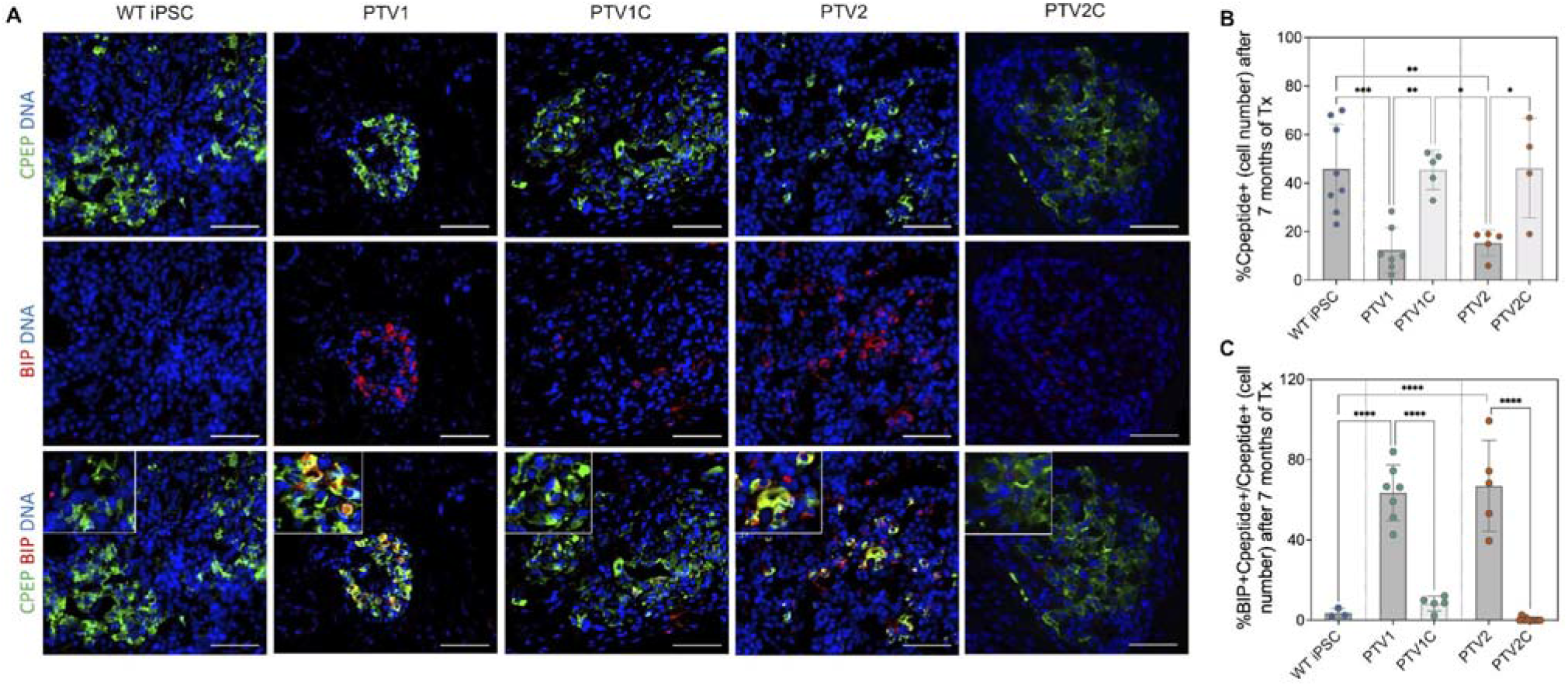
Increased expression of ER-stress markers grafts with PTV mutations. **(A)** Immunohistochemistry showing expression of ER stress marker-BIP in WT iPSC, PTV1, PTV1C, PTV2, and PTV2C SC-islets excised at 7 months post-transplantation. Scale bar: 50μm. **(B)** Quantification of percentage of C-peptide+ cells. **(C)** Quantification of colocalization of C-peptide+ and BIP+.The C-peptide antibody used detects both C-peptide and aa 33-63 of proinsulin. n=3-4 independent transplanted mice per cellular genotype. Data were quantified by cell numbers and presented as mean ± SEM. One-way ANOVA test with *P<0.05, **P < 0.01, ***P < 0.001, ****P < 0.0001.

Introduction of the PTV1 mutation in Mel1 cells also reduced the number of C-peptide positive cells in grafts, and increased the BIP+C-peptide+ population from zero to 80% (**Figure S6A, C**). These results (**Figure 4B)** indicate significant beta cell loss resulting from the presence of the mutant *INS* alleles.

Supporting the possibility that apoptosis contributes to at least some of the beta cell loss in mutant beta cells (Laybutt et al., 2007), we observed a four-fold increase in TUNEL-positive, C-peptide-producing cells in PTV1 cells compared to corrected PTV1C cells after 27 days of *in vitro* differentiation (**Figure S6D, E**). A similar trend was seen in cells with the PTV2 mutation compared to the gene-corrected isogenic control cells, but this difference did not achieve statistical significance (**Figure S6D, E**). However, even for the PTV1 cells, apoptosis was nearly undetectable in transplanted beta cells at seven months after in vivo engraftment (**Figure S6F**), raising the question of whether other mechanisms might also contribute to beta cell loss.

### Beta cells with INS mutation lost their beta cell identity after transplantation as a consequence of dedifferentiation

We immunostained stem cell-derived beta cell clusters for the presence of the alpha cell marker, glucagon, and assessed the ratio of C-peptide+ cells to Glucagon+ cells before and after 7 months of transplantation. Both controls (WT-iPSC) and corrected isogenic controls had similar percentage of C-peptide positive cells as a fraction of total cells prior to transplantation (**Figure 5A, C**). There was a ∼70% reduction of c-peptide positive beta-like cells in PTV1 and PTV2 mutant grafts at 7 months post-transplantation (**Figure 5B, C**). The ratio of proinsulin or C-peptide-positive cells / Glucagon-positive cells was approximately 1.6 in isogenic corrected and WT groups both before and after transplantation. In contrast, PTV1 and PTV2 grafts exhibited a decline in this ratio: decreasing from 1.7 (PTV1) and 1.5 (PTV2) before transplantation, to 0.5 (PTV1, P=0.002) and 0.4 (PTV2, P=0.012) at 7 months post-transplantation (**Figure 5A, B** and **D**). We observed the most significant reduction in this ratio in Mel1-PTV1 grafts, decreasing from 1.7 to 0.1 after 7 months of transplantation (P<0.0001) (**Figure S7A, B**).

**Figure 5.**
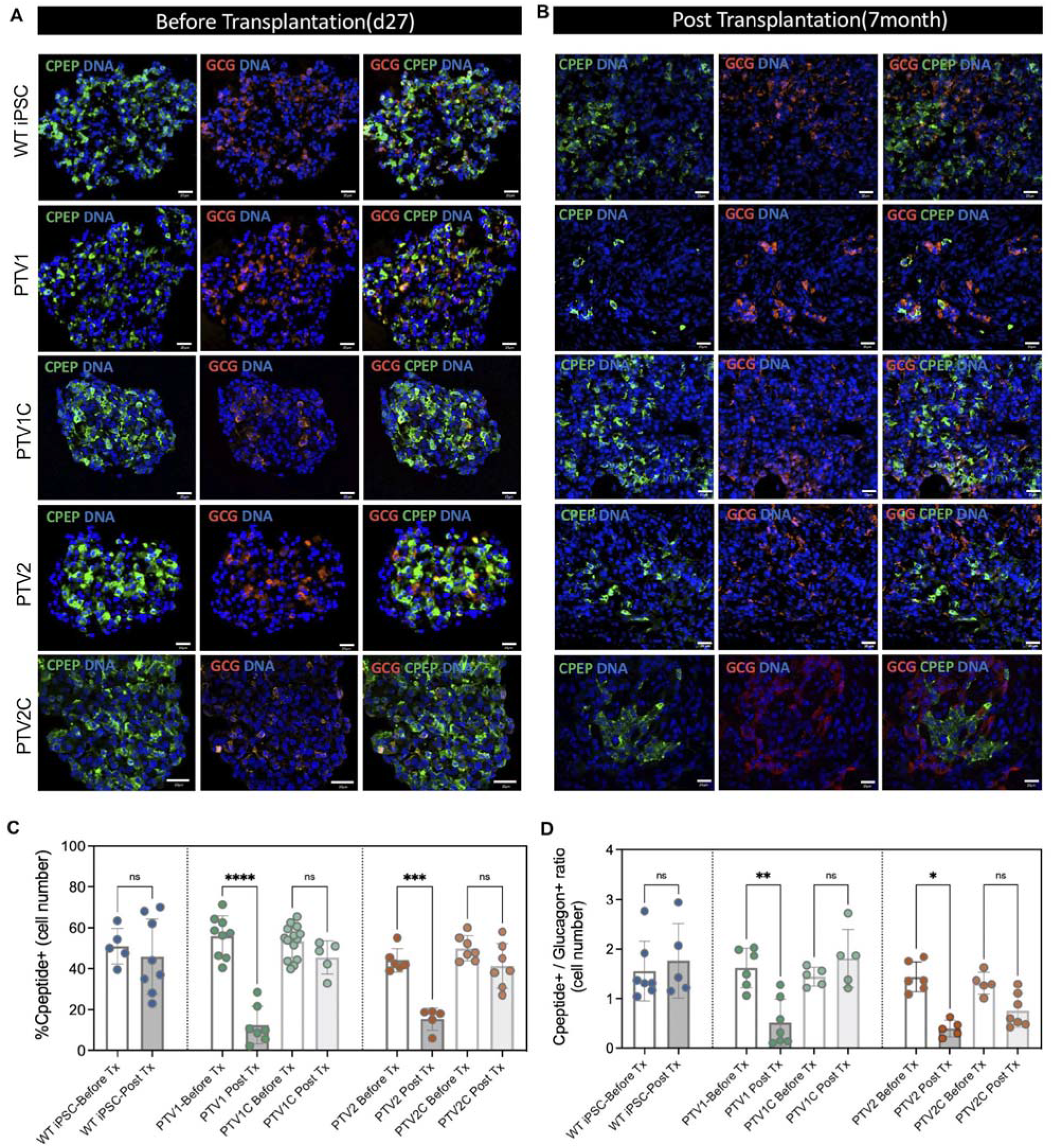
Loss of Insulin producing cells in PTV SC-islets engrafted to NSG mice. Immunohistochemistry for beta cell marker C-peptide and alpha cell marker Glucagon at d27 of in vitro differentiation. Scale bar: 20μm. **(B)** Immunohistochemistry for beta cell marker C-peptide and alpha cell marker Glucagon at 7 months post transplantation. Scale bar: 20μm. **(C)** Comparisons of percentage of C-peptide positive cells (%C-peptide+/Hoechest+) before(d27) and after transplantation (7months). n=3-4 independent transplanted mice per cell genotype. Two-way ANOVA. **(D)** Ratio of insulin-producing cell number to Glucagon-producing cell number in vitro (27d) and in vivo (7months). The C-peptide antibody used detects both C-peptide and aa 33-63 of proinsulin. Cell numbers are presented as mean ± SEM. Two-way ANOVA test with *P<0.05, **P < 0.01, ***P < 0.001, ****P < 0.0001.

Specific factors can trigger loss (dedifferentiation) of mature beta cell identity. These include: oxidative stress (Kitamura et al., 2005; Tanaka et al., 1999), ER stress (Chen et al., 2022), inflammation (Nordmann et al., 2017) and hypoxia (Puri et al., 2013). We performed immunoreactivity assays for ALDH1A3 (**Figure 6A and Figure S8A**), which is upregulated in diabetic beta cell failure (Cinti et al., 2016; Talchai et al., 2012). The mean percentage of ALDH1A3-positive cells per block was elevated more than 3-fold in PTV grafts compared with controls (P=0.0012 for Mel1-PTV1 vs Mel1-WT) and isogenic corrected controls (P< 0.001 for PTV1 vs PTV1C and P=0.037 for PTV2 vs PTV2C) (**Figure 6B and Figure S8B**). In PTV1 grafts, more than 30% of ALDH1A3+ cells expressed NKX6.1, a beta cell specification transcription factor; the percentage of ALDH1A3+NKX6.1+ cells in PTV1 grafts was about 17 times higher than in isogenic control grafts (P=0.016) (**Figure 6C**). In PTV2 patient cells, total ALDH1A3+ cells and ALDH1A3+ cells co-expressing NKX6.1 were also reduced after gene correction, although the difference was less significant (P= 0.036 in ALDH1A3+ cells and P=0.038 in ALDH1A3+NKX6.1+ cells) (**Figure 6C**). To further examine whether insulin mutations are associated with molecular stigmata of de-differentiation, we examined iMel1-PTV1 grafts. Over 20% of ALDH1A3+ cells co-expressed NKX6.1, and the percentage of these double-positive cells were 4-fold higher than in Mel1-WT grafts (P<0.0001) (**Figure S8C**). These findings are consistent with the inference that ALDH1A3 positive cells that are also positive for NKX6.1 are de-differentiated beta cells (Cinti et al., 2016). Of note is that the ALDH1A3+NKX6.1+ double-positive populations were no longer expressing insulin (**Figure 6A and Figure S8A**).

**Figure 6.**
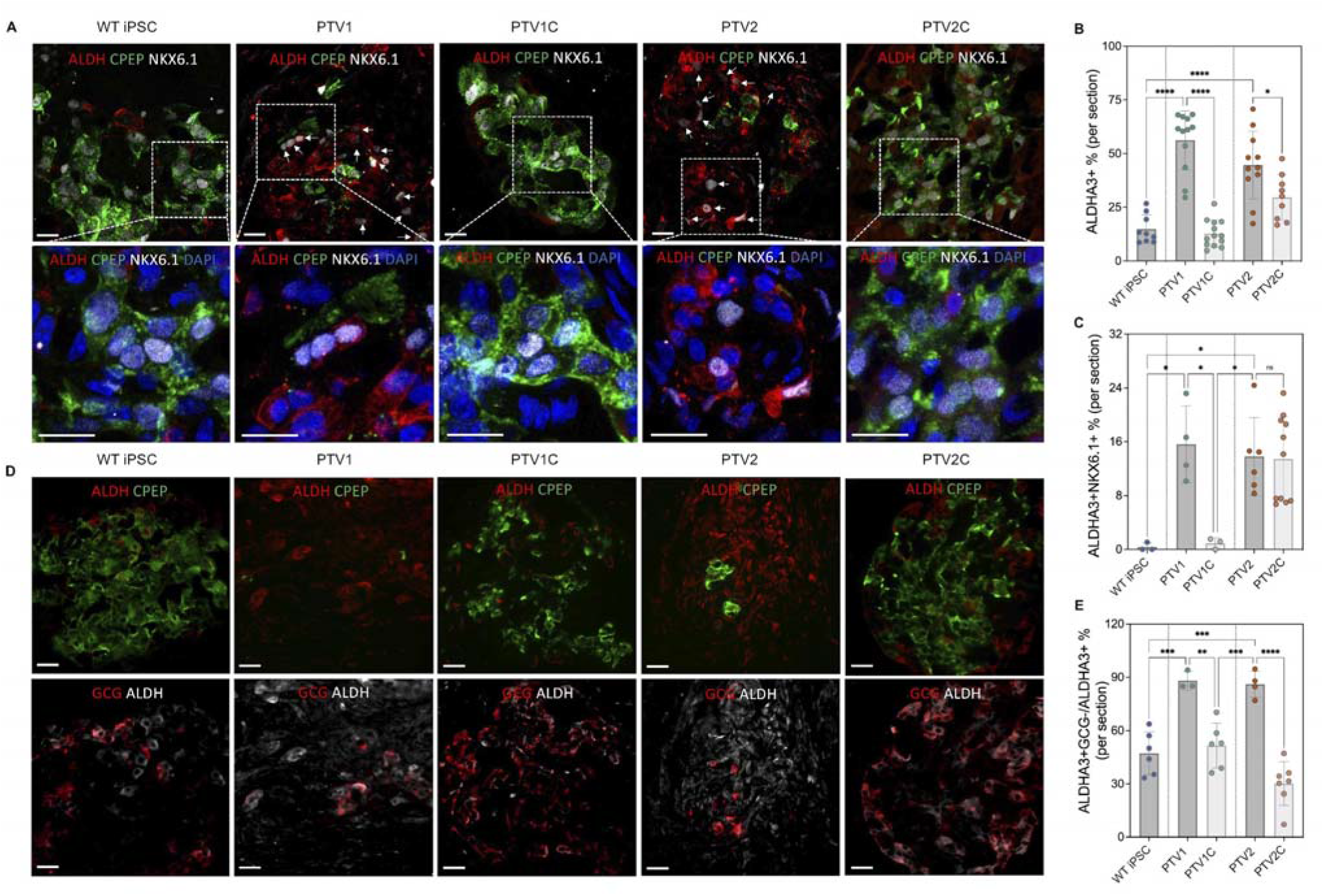
Increased beta cell dedifferentiation in stem cell-derived islet-like grafts carrying PTV mutations. **(A)** Immunohistochemistry for colocalization of dedifferentiation marker ALDH1A3 (ALDH) with beta cell marker C-peptide, NKX6.1 in 7-month-old grafts. Scale bar: 10μm. **(B)** Percentage of ALDH1A3 positive cells. **(C)** Percentage of NKX6.1 positive cells co-expressing ALDH1A3, indicating dedifferentiated beta cells. **(D)** Immunohistochemistry for colocalization of dedifferentiation marker ALDH1A3 (ALDH) with C-peptide and alpha cell marker Glucagon in 7-month-old grafts. The C-peptide antibody used detects both C-peptide and aa 33-63 of proinsulin. Scale bar: 20μm. **(E)** Quantitation of GCG-negative ALDH1A3 expressing cells. Each data point represents an independent frozen section randomly selected from 3-4 independent transplanted mice per cell genotype. Cell numbers are presented as mean ± SEM. One-way ANOVA test with *P<0.05, **P < 0.01, ***P < 0.001, ****P < 0.0001.

ALDH1A3 is normally expressed in alpha cells, which are increased in T2D. Glucagon negative alpha cells are thought to be a consequence of beta cell de-differentiation (Cinti et al., 2016). The percentage of GCG-negative ALDHA3-expressing cells, representing a progenitor cell-like stage, was significantly increased in PTV grafts compared with their isogenic controls in both patient genotypes (P=0.0013 for PTV1 vs PTV1C, P<0.0001 for PTV2 vs PTV2C) (**Figure 6D, E**). A difference was also seen in Mel1-PTV1 grafts when compared with Mel1-Wt controls, though it did not reach statistical significance (**Figure S8D, E**).

Collectively, these results suggest that beta cells subjected to ER stress by misfolded insulin molecules exhibit heightened dedifferentiation, characterized by increased ALDH1A3 expression and loss of insulin expression.

## Discussion

We used patient-derived iPSCs as well as mutated human ESCs bearing two distinct clinically dominant *INS* mutations to identify genetically dominant derangements in proinsulin processing, proinsulin folding, ER trafficking and insulin production, associated with increased ER stress, beta cell dedifferentiation and death. These molecular-cellular phenotypes are consistent with the early onset insulin deficient phenotype of patients with these mutations.

Consistent with clinical data, stem cell-derived beta cell grafts harboring these specific mutations - in comparison to isogenic controls - produced less human insulin and showed reduced insulin-expressing cell numbers after 7 months of transplantation. This decrease in the number of insulin-producing cells was due in part to cellular de-differentiation with loss of insulin protein production. From these findings, we infer that beta cells carrying the implicated neonatal diabetes *INS* mutations are subjected to proteotoxic misfolding of insulin precursors with attendant elevated ER stress and impaired insulin secretion. In addition, apoptosis, and beta cell de-differentiation reduce the number of insulin secreting cells. Thus, the clinical phenotype of severe insulin insufficiency results from convergent molecular and cellular events compromising the production of insulin.

Proinsulin misfolding has been proposed as a mechanism in PNDM (Elizabeth Jane Besser et al., 2016; Shankar et al., 2013). These patients frequently segregate for heterozygous missense *INS* mutations leading to early onset (generally within the 1st 6 months of life) of permanent insulin deficiency (Dahl and Kumar, 2020; Iafusco et al., 2002). Heterozygosity for null mutations of *INS* predisposes to diabetes, but of later onset and presumably in individuals with an enabling genetic background(Carmody et al., 2015; Raile et al., 2011). Thus, dominantly inherited *INS* mutations implicated in early onset insulin-dependent diabetes must have consequences for insulin processing and/or the viability of cells expressing the mutant peptide. This inference is supported by human clinical phenotypes as well as animal models including mouse and transgenic pigs (Blutke et al., 2017; Oyadomari et al., 2002; Renner et al., 2013). In Akita mice expressing 3 copies of wild type *INS* (two *INS*1 and one *INS*2) and 1 copy of a misfolding *INS* mutation (*INS2*^+/C96Y^), mutated proinsulin protein accounting for 36% of total translated proinsulin products was misfolded to higher-molecular-weight forms and that were associated with activation of the UPR (Liu et al., 2010). We observed dimerized and trimerized proinsulin, as well as an elevated ratio of each higher-molecular-weight isoform to its monomer in cells expressing L^B15^Y^B16^delinsH (PTV1) or Y^B26^C (PTV2) heterozygous mutations. These mutations interfere with the cleavage and processing of proinsulin, and prohibit the formation of disulfide bonds in mature insulin (Liu et al., 2018; Qiao et al., 2001). A dominant-negative effect of the mutant insulin on processing of the wild type allele has been demonstrated (Liu et al., 2007). This dominant effect *in vitro* in sc-beta cells, which resemble fetal-like beta cells (Hrvatin et al., 2014). This effect reduces, but does not eliminate all insulin secretion. Additional cellular consequences account for the long-term decline of insulin production after birth and diabetes. Heterozygous *INS* mutations may also cause diabetes with later onset in childhood, mimicking type 1 diabetes (Bonfanti et al., 2009; Yu et al., 2019). In these instances, C-peptide is usually detectable at the time of diagnosis and persists in gradual decline, suggesting that cell death and/or de-differentiation are contributing to the hypoinsulinemic phenotype. These long-term consequences are more accurately modeled by grafting of beta cells in mice (this study, and González et al., 2022).

We observed severe reductions of insulin-producing cells in both L^B15^Y^B16^delinsH and Y^B26^C islet-like grafts at 7-months post transplantation. The majority of the surviving insulin-producing cells showed evidence of increased ER stress characterized by elevation of canonical molecular markers. However, given the impacts of L^B15^Y^B16^delinsH and Y^B26^C mutations on *INS* folding we identified *ex vivo*, these findings are not surprising. Beta-cell apoptosis has been proposed as a main consequence of insulin misfolding and ER stress in many types of diabetes, including in T2D (Laybutt et al., 2007; Oyadomari et al., 2002), MODY (Wang et al., 1999) and permanent neonatal diabetes (PND) (Colombo et al., 2008; Støy et al., 2007). The L^B15^Y^B16^delinsH mutation in patient PTV1 has been previously shown to cause apoptosis in cell-based assays (Colombo et al., 2008). Similarly, we detected increased apoptosis by TUNEL staining in stem cell derived beta cells *in vitro*. In contrast, we saw little apoptosis in transplanted beta-cells over a 7-month period. TUNEL may have missed a transient *in vitro* signal and *in vivo* adaptations may have mitigated longer term effects of the mutations. Compensatory reductions in (pro)insulin biosynthesis can reduce ER stress (Ron, 2002). The Y^B26^C mutation in PTV2 patients seems to have a greater effect on insulin synthesis and beta cell dedifferentiation than apoptosis. PNDM mutation *INS* C96R impacts a crucial cysteine residue responsible for the inter-chain disulfide bonds A7-B7; higher levels of ER stress were observed both *in vitro* and *in vivo* models of this mutation (Balboa et al., 2018). These findings are consistent with the activated unfolded protein response (UPR) noted in our study. But no increase in apoptosis was noted at any stage of differentiation in *INS* C96R cells (Balboa et al., 2018). Apoptosis may not be the primary cause of loss of beta cell function in PNDM caused by insulin mutations.

Here we have identified beta cell dedifferentiation as an important possible mechanism for loss of beta cell function in PNDM due to dominant/negative *INS* mutations as reflected in the presence of indicative molecular signatures: ALDH1A3+NKX6.1+ and ALDH1A3+ GCG-cells in SC-islet-like grafts after long term *in vivo* maturation. In agreement with a recent study showing a significant increase of Aldh1a3+ cells in islets of Akita mice at 5 months after birth, we found a distinct increase of Aldh1a3 expression in mutant transplants after months of *in vivo* maturation (Kitakaze et al., 2021). Upregulation of ALDH1A3 has been confirmed as a signature of dedifferentiation in human T2D diabetes islets, but has not been previously shown to occur in *INS* mutated human islets, or associated with ER stress caused by protein misfolding. Stem cell models of beta cell dysfunction involving ER stress have been previously reported (Balboa et al., 2018; Shang et al., 2014), but were not examined for evidence of de-differentiation.

Our data implicate beta cell de-differentiation as a proximate mechanism for the beta cell failure characteristic of PNDM resulting from dominant-negative *INS* mutations. Hyperglycemia-induced ER stress is characteristic of T2D (Laybutt et al., 2007), and likely contributes to beta cell de-differentiation. Other genetic forms of neonatal diabetes due to explicit ER stress, including Wolfram and Wolcott-Rallison syndromes (Stone et al., 2021) may also be associated with beta cell de-differentiation. That ER stress (perhaps depending on the duration and intensity) can have various outcomes, including beneficial ones, such as cell proliferation (Sharma et al., 2015; Snyder et al., 2021). In the case of insulin folding mutation, lasting stress has the consequence of de-differentiation.

Our results have implications regarding the biology of diabetes at several levels. They are consistent with recent suggestions that therapeutic Inhibition of ALDH1A3 might be useful to mitigate loss of functional beta cells in diabetes (Son et al., 2023). They also point to the likely clinical utility of isogenic stem cell lines in the creation of genetically “corrected” beta cells for transplantation in patients with monogenic etiologies of diabetes. However, such therapeutic advances will also require parallel improvements in iPSC reprogramming and quality control, as the differentiation potential of iPSCs to insulin producing cells often remains very variable (this study &Sui et al., 2018). And they again emphasize the utility of human stem cell-derived beta cells - and their *in vitro* and *in vivo* analyses - in understanding the molecular physiology of specific genetic derangements leading to clinical diabetes (González et al., 2022; Shang et al., 2014).

## Materials and Methods

### Patients and cell lines

This study includes 10 human cell lines: a. 4 human embryonic stem cell lines (Mel1-WT: *INS* ^+/GFP^, Mel1-PTV1: *INS* ^PTV1/GFP^, Mel1-PTV2: *INS* ^PTV1/GFP^, Mel1-INS KO: *INS* ^-/GFP^). Mel1-WT was generated previously (Micallef et al., 2012); the other three were gene-edited using CRISPR/Cas9. b. 6 human pluripotent stem cell lines: (WT-iPSC: *INS* ^+/+^, PTV1: *INS* ^PTV1/+^, PTV2: *INS* ^PTV2/+^, PTV1C1: *INS* ^+/+^, PTV1C2: *INS* ^+/+^, PTV2C: *INS* ^+/+^). WT-iPSC was derived from a healthy donor (cell line 1159) (Yamada et al., 2014); PTV1 and PTV2 were reprogrammed from 2 PNDM patients as described in Results. 2 isogenic controls denominated PTV1C1 and PTV1C2 were corrected from PTV1 and another isogenic control PTV2C was corrected from PTV2. iPSCs derived from healthy or diabetic donors were reprogrammed from fibroblasts using mRNA reprogramming kit (Ma et al., 2018). Additional details including cell lines, gRNAs, ssDNA templates and PCR primers are provided in **Table S2** and **Table S3**.

Cell lines were karyotyped by G band karyotyping by Cell Line Genetics. Karyotyping was also performed using low pass whole genome sequencing. Libraries were prepared using the Illumina DNA prep kit (catalog 20060059). Sequencing was performed using Illumina HiSeq 2500 with 2×150bp configuration. Sequenced samples were analyzed using bwa aligner (v.0.7.17), samtools (v.1.11), and R package QDNAseq (v1.26.0.) Karyotypes were inspected using quantification of read numbers with a bin size of 500kb.

Human fibroblasts were obtained after informed consent of the legal guardians of the patient at Salesi Hospital. Samples were de-identified and numbered PTV1, PTV2. Genetic analysis and use of patient samples in research was done with IRB approval at the University of Rome Tor Vergata. Work with human pluripotent stem cell lines was reviewed and approved by the Columbia University Embryonic Stem Cell Committee.

### INS variants correction by CRISPR/Cas9

CRISPR/Cas9 gene editing system was employed to correct 2 pathogenic variants of *INS* (PTV1: L^B15^Y^B16^delinsH, PTV2: Y^B26^C) and to generate 3 corrected cell lines (PTV1C1, PTV1C2 and PTV2C). Guide RNAs against *INS* loci adjacent to the mutations were designed using the online tool (https://www.idtdna.com/pages/tools/alt-r-crispr-hdr-design-tool) and synthesized by IDT (Integrated DNA Technologies). gRNAs were then assembled in a backbone vector (gBlock) by PCR for further functional expression using the method of Gibson (Gibson et al., 2009). gBlocks, correction template composed of single strand DNA (IDT) and Cas9-GFP were then transfected into human iPSCs for *INS* mutation correction using embryonic stem cell Nucleofector Kit (VVPH-5012, Lonza). Transfected cells were cultivated in StemFlex medium (catalog #A3349401, Thermo Fisher Scientific) supplemented with 10 mmol/L ROCK inhibitor-Y26732 (catalog #S1049; Selleckchem). After 48 hours of proliferation, GFP positive cells were sorted by flow cytometry facilities in Columbia stem cell core and seeded onto GelTrex (catalog A1413302, Thermo Fisher Scientific) coated plates. ROCK inhibitor was removed after 24 hours and the medium was refreshed every other day until 2 weeks later. Each clone was picked up and analyzed by PCR and sanger sequencing to select corrected *INS* clones. Additional details including cell lines, gRNAs, ssDNA templates and PCR primers are provided in **Table S2 and Table S3**.

### Cell culture and SC-derived beta-cell differentiation

All iPSCs and ESCs employed in this paper were seeded on GelTrex-coated plates, cultured in StemFlex medium and passaged with TrypLE Express (catalog #12605036; Life Technologies) every 3-5 days. Undifferentiated cells were then cultivated following a step-wise differentiation protocol previously established by our lab (Sui et al., 2018b); graphics illustrating differentiation stages are provided in Figure S1A. Cells were aggregated into clusters at Pancreatic Progenitor stage and were collected for further analysis within pancreatic beta-cell stage on d27. Images showing GFP-positive insulin-expressing cells were captured by an OLYMPUS IX73 fluorescent microscope.

### Flow cytometry analysis

The SC-derived islet-like clusters were dissociated into single cells on d27 of differentiation using TrypLE Express. Dispersed cells were suspended in 5% FBS-containing PBS buffer for further use. Cells were then fixed in 4% paraformaldehyde (PFA) for 10 min followed by permeabilization at 20°C with cold methanol for 10 min. Primary antibodies were incubated with cells for 1 hour at 4°C with a dilution of 1:100 in 0.5%BSA. Secondary antibodies were incubated with cells at a dilution of 1:500 for 1 hour at room temperature. Cells were washed twice between every step by 5% FBS-containing PBS buffer. Lastly, collected cells were filtered into a Falcon round-bottom 12 × 15mm tube through the cell strainer cap (catalog #352235; Corning) and analyzed by flow cytometer in Columbia stem cell core. All staining was performed at 4°C. Data analysis was performed on FlowJo Software. Refer to **Table S4** for additional information of primary antibodies and **Table S5** for secondary antibodies.

### Immunohistochemistry

SC-derived islet-like clusters were collected at day 27 of culture and fixed with 4% PFA at room temperature for 10-20 minutes. Grafts were retrieved from the NSG mice post 7 months of transplantation, fixed with 4% PFA at room temperature for 1 hour. The next steps were performed according to our formerly published method (Sui et al., 2018b). Briefly, fixed clusters and grafts were washed twice by PBS and incubated with 30% sucrose overnight at 4°C to dehydrate and precipitate clusters. Aggregated clusters were transferred into a cryomold and immersed with O.C.T. medium over dry ice for cryosection. The frozen blocks were cut into 5-μm sections sections for further immunostaining. Primary antibodies used with this assay are listed in **Table S4**, and secondary antibodies are listed in **Table S5**. Images were taken with an OLYMPUS IX73 fluorescent microscope or a ZEISS LSM 710 confocal microscope. Frozen sections stained with the same markers were performed simultaneously and imaged with same parameters settings to ensure reliable quantification and comparison. Imaging processing and later quantification were performed blind, without labels using Fiji Software. Cell numbers were counted manually to calculate the positive or negative cell populations in all immunostaining sections. Apoptosis analysis was carried out with TUNEL assay. Frozen sections were processed with TUNEL apoptosis detection kit-CF594(catalog #30064, Biotium) following the manufacturer’s instructions.

### Western blotting

SC-derived islet-like clusters at d27 were collected into pre-chilled 1.5ml tubes and lysed with RIPA at 4°C for 30 mins. Clusters were vortexed every 5min to ensure sufficient lysis. Supernatants containing total protein were retrieved after centrifuge. Samples were resolved by SDS in 4–12% Bis–Tris NuPAGE gels under either non-reducing or reducing conditions and electrotransferred to nitrocellulose. Development of immunoblots used enhanced chemiluminescence, captured with a Fotodyne gel imager, quantified using Fiji software. Of note, anti-KDEL antibodies were used to detect both Grp94 and BIP. Additional details including primary and secondary antibodies are provided in **Table S4 and Table S5**.

### Metabolic Labeling and Immunoprecipitation

The pulse-chase experiment was performed as described previously (Haataja et al., 2016). Briefly, SC-derived islet-like clusters at d27 were dissociated to ∼2-4 million cells and then labeled with ^35^S-Cys/Met by incubation with cysteine/methionine-absent media for 30 min and defined pulse chase times. Labeled cells were washed with PBS containing 20 mmol/L N-ethyl maleimide (NEM) and then lysed in radioimmunoprecipitation assay buffer (25 mmol/L Tris, pH 7.5; 1% Triton X-100; 0.2% deoxycholic acid; 0.1% SDS; 10 mmol/L EDTA; and 100 mmol/L NaCl) with 2 mmol/L NEM and protease inhibitor. Centrifuged cell lysates, normalized to trichloroacetic acid precipitable counts and precleared with zysorbin, were immunoprecipitated and probed with anti-proinsulin and anti-insulin (B-chain) antibodies overnight at 4°C. Immunoprecipitates were analyzed by nonreducing Tris–tricine–urea– SDS-PAGE, with phosphorimaging.

### ELISA assays

Human proinsulin and insulin levels measured from cell content using human Proinsulin ELISA kit (Catalog #10-1118-01, Mercodia) and human Insulin ELISA kit (Catalog #10-1113-01, Mercodia). Human C-peptide level in mice serum sample were detected by Ultra-sensitive C-peptide ELISA kit (Catalog #10-1141-01, Mercodia). Mouse C-peptide levels were measured from plasma samples by using mouse C-peptide ELISA kit (Catalog #90050, Crystal Chem). All procedures were performed according to their manufacturing instructions.

### Transplantation to NSG mice of SC-derived islet-like clusters and their in vivo assay

8-week-old male NOD-SCID-gamma (NSG) mice were purchased from Jackson Laboratories (catalog #005557) and housed at Columbia University Medical Center animal facility. Intra-leg quadriceps muscle transplantations were performed in 8 to10-week-old mice with approximately 2 million beta cells per mouse in accordance with published methods (González et al., 2022). Briefly, SC-derived islet-like clusters were collected at d27 into a 1.5ml pre-chilled tube with 50 μL Matrigel. Mice were anesthetized with isoflurane. Clusters mixed with gel were injected into the quadriceps using a 21-gauge needle. Fed state human C-peptide levels were detected and recorded (@ 4-5pm) at 2 weeks post engraftment and every subsequent month. Glucose-stimulated insulin secretion was performed by fasting overnight (15-17h) and 2g/kg D-glucose intraperitoneal injection. Blood was collected from the tail vein into heparin-coated tubes at fed state, fasting state (0min) and 30min post injection (30min). Whole blood glucose was also measured by glucometer at each time point. Supernatant plasma was separated through centrifugation at 2000 g for 15 min at 4 °C for human and mouse C-peptide measurement.

### Statistical analysis

Statistical analysis was performed with Prism software (GraphPad Prism 9, GraphPad Software, Inc.). Differences between experimental groups were tested by 2-tailed unpaired or paired t test and by one-way or two-way ANOVA followed by Tukey’s multiple-comparison test and presented as mean ± standard error of mean (SEM). The differences observed were considered statistically significant at the 5% level and annotated: *P < 0.05, **P < 0.01, ***P < 0.001, ****P < 0.0001.

### Manuscript preparation

This manuscript was prepared in part using solar electricity collected with a V250 Voltaic Systems solar panel and external laptop battery between sea level in NYC and 17,200 feet in Denali National Park in the United States. Figure schematics were prepared using Biorender.

## Author contributions

LS, YZ, CUN, FB, PA and DE conceived, discussed and designed the study. LS and YZ carried out PTV1 patient-derived iPSCs relative experiments, YZ performed most PTV2 patient-derived iPSCs and Mel1-ESCs relative experiments, combined and analyzed all data, and wrote the manuscript with contributions from LS and DE, as well as from LH, PA and FB. LH and PA performed Western Blot experiments, design and interpretation. QD performed mice transplantation experiments, *in vivo* measurement and immunostaining for PTV2C grafts and Mel1-PTV1 grafts. YY assisted with generating mutated Mel1 cell lines, performed relative differentiation experiments and corresponding immunohistochemistry experiments. SX performed library preparation and karyotype analysis using sequencing read quantification. RV performed iPS reprogramming. RL contributed supervision. CUN conducted initial studies on iPS derivation, correction and differentiation. FB coordinated human subject research. DE provided support, and supervision throughout the study.

## Acknowledgments

This research was supported by the American Diabetes Association (grant #1-16-ICTS-029) and the NYSTEM IDEA award # C029552, Leona and Harry Helmsley Charitable Trust, Helmsley Trust Diabetes Cell Repository, the Juvenile Diabetes Research Foundation (JDRF), the Naomi Berrie Foundation program for Cellular Therapies of Diabetes, F.B. was supported by and the Italian Ministry of Health (project PE-2011-02350284). The contributions of P.A. and L.H. were supported by NIH R01 DK48280. These studies used the resources of the Herbert Irving Comprehensive Cancer Center Flow Cytometry Shared Resources funded in part through Center Grant P30CA013696, and the Diabetes Research Center Flow Core Facility funded in part through DRC Center Grant (5P30DK063608) and the MBMG Core in the New York Nutrition and Obesity Research Center (5P30DK026687). We thank Charles LeDuc for help with mouse studies, Rudolph Leibel for helpful input and critical reading of the manuscript, and Robin Goland for helpful discussions on beta cell replacement therapies. We appreciate Stefano Zucchini and Federica Ortolani for their clinical contributions to PTV1 and PTV2 patients.

## Reference

1. Abdelalim, E.M., 2020. Modeling different types of diabetes using human pluripotent stem cells. Cell. Mol. Life Sci. 2020 786 78, 2459–2483. 10.1007/S00018-020-03710-9

2. Alam, M., Arunagiri, A., Haataja, L., Torres, M., Larkin, D., Kappler, J., Jin, N., Arvan, P., 2021. Predisposition to Proinsulin Misfolding as a Genetic Risk to Diet-Induced Diabetes. Diabetes 70, 2580–2594. 10.2337/DB21-0422

3. Arunagiri, A., Haataja, L., Pottekat, A., Pamenan, F., Kim, S., Zeltser, L.M., Paton, A.W., Paton, J.C., Tsai, B., Ansari, P.I., Kaufman, R.J., Liu, M., Arvan, P., 2019. Proinsulin misfolding is an early event in the progression to type 2 diabetes. Elife 8. 10.7554/ELIFE.44532

4. Balboa, D., Saarimäki-Vire, J., Borshagovski, D., Survila, M., Lindholm, P., Galli, E., Eurola, S., Ustinov, J., Grym, H., Huopio, H., Partanen, J., Wartiovaara, K., Otonkoski, T., 2018. Insulin mutations impair beta-cell development in a patient-derived iPSC model of neonatal diabetes. Elife 7, 1–35. 10.7554/eLife.38519

5. Blutke, A., Renner, S., Flenkenthaler, F., Backman, M., Haesner, S., Kemter, E., Ländström, E., Braun-Reichhart, C., Albl, B., Streckel, E., Rathkolb, B., Prehn, C., Palladini, A., Grzybek, M., Krebs, S., Bauersachs, S., Bähr, A., Brühschwein, A., Deeg, C.A., De Monte, E., Dmochewitz, M., Eberle, C., Emrich, D., Fux, R., Groth, F., Gumbert, S., Heitmann, A., Hinrichs, A., Keßler, B., Kurome, M., Leipig-Rudolph, M., Matiasek, K., Öztürk, H., Otzdorff, C., Reichenbach, M., Reichenbach, H.D., Rieger, A., Rieseberg, B., Rosati, M., Saucedo, M.N., Schleicher, A., Schneider, M.R., Simmet, K., Steinmetz, J., Übel, N., Zehetmaier, P., Jung, A., Adamski, J., Coskun, Ü., Hrabě de Angelis, M., Simmet, C., Ritzmann, M., Meyer-Lindenberg, A., Blum, H., Arnold, G.J., Fröhlich, T., Wanke, R., Wolf, E., 2017. The Munich MIDY Pig Biobank – A unique resource for studying organ crosstalk in diabetes. Mol. Metab. 6, 931–940. 10.1016/J.MOLMET.2017.06.004

6. Bonfanti, R., Colombo, C., Nocerino, V., Massa, O., Lampasona, V., Iafusco, D., Viscardi, M., Chiumello, G., Meschi, F., Barbetti, F., 2009. Insulin gene mutations as cause of diabetes in children negative for five type 1 diabetes autoantibodies. Diabetes Care 32, 123–125. 10.2337/DC08-0783

7. Brozzi, F., Eizirik, D.L., 2016. ER stress and the decline and fall of pancreatic beta cells in type 1 diabetes. https://mc.manuscriptcentral.com/ujms 121, 133–139. 10.3109/03009734.2015.1135217

8. Brusko, T.M., Russ, H.A., Stabler, C.L., 2021. Strategies for durable β cell replacement in type 1 diabetes. Science 373, 516–522. 10.1126/SCIENCE.ABH1657

9. Carmody, D., Park, S.Y., Ye, H., Perrone, M.E., Alkorta-Aranburu, G., Highland, H.M., Hanis, C.L., Philipson, L.H., Bell, G.I., Greeley, S.A.W., 2015. Continued lessons from the INS gene: an intronic mutation causing diabetes through a novel mechanism. J. Med. Genet. 52, 612–616. 10.1136/JMEDGENET-2015-103220

10. Chen, C.W., Guan, B.J., Alzahrani, M.R., Gao, Z., Gao, L., Bracey, S., Wu, J., Mbow, C.A., Jobava, R., Haataja, L., Zalavadia, A.H., Schaffer, A.E., Lee, H., LaFramboise, T., Bederman, I., Arvan, P., Mathews, C.E., Gerling, I.C., Kaestner, K.H., Tirosh, B., Engin, F., Hatzoglou, M., 2022. Adaptation to chronic ER stress enforces pancreatic β-cell plasticity. Nat. Commun. 2022 131 13, 1–18. 10.1038/s41467-022-32425-7

11. Cinti, F., Bouchi, R., Kim-Muller, J.Y., Ohmura, Y., Sandoval, P.R., Masini, M., Marselli, L., Suleiman, M., Ratner, L.E., Marchetti, P., Accili, D., 2016. Evidence of β-Cell Dedifferentiation in Human Type 2 Diabetes. J. Clin. Endocrinol. Metab. 101, 1044–1054. 10.1210/JC.2015-2860

12. Colombo, C., Porzio, O., Liu, M., Massa, O., Vasta, M., Salardi, S., Beccaria, L., Monciotti, C., Toni, S., Pedersen, O., Hansen, T., Federici, L., Pesavento, R., Cadario, F., Federici, G., Ghirri, P., Arvan, P., Iafusco, D., Barbetti, F., 2008. Seven mutations in the human insulin gene linked to permanent neonatal/infancy-onset diabetes mellitus. J. Clin. Invest. 118, 2148. 10.1172/JCI33777

13. Dahl, A., Kumar, S., 2020. Recent Advances in Neonatal Diabetes. Diabetes, Metab. Syndr. Obes. Targets Ther. 13, 355. 10.2147/DMSO.S198932

14. De Franco, E., Flanagan, S.E., Houghton, J.A.L., Allen, H.L., MacKay, D.J.G., Temple, I.K., Ellard, S., Hattersley, A.T., 2015. The effect of early, comprehensive genomic testing on clinical care in neonatal diabetes: an international cohort study. Lancet (London, England) 386, 957–963. 10.1016/S0140-6736(15)60098-8

15. Edghill, E.L., Dix, R.J., Flanagan, S.E., Bingley, P.J., Hattersley, A.T., Ellard, S., Gillespie, K.M., 2006. HLA Genotyping Supports a Nonautoimmune Etiology in Patients Diagnosed With Diabetes Under the Age of 6 Months. Diabetes 55, 1895–1898. 10.2337/db06-0094

16. Edghill, E.L., Flanagan, S.E., Patch, A.M., Boustred, C., Parrish, A., Shields, B., Shepherd, M.H., Hussain, K., Kapoor, R.R., Malecki, M., MacDonald, M.J., Støy, J., Steiner, D.F., Philipson, L.H., Bell, G.I., Hattersley, A.T., Ellard, S., 2008. Insulin mutation screening in 1,044 patients with diabetes mutations in the INS gene are a common cause of neonatal diabetes but a rare cause of diabetes diagnosed in childhood or adulthood. Diabetes 57, 1034–1042. 10.2337/db07-1405

17. Elizabeth Jane Besser, R., Mrcpch, M., Elizabeth Flanagan, S., Gordon Mackay, J., Karen Temple, I., Frcp, Mbc., Helen Shepherd, M., Maureen Shields, B., Ellard, S., Tym Hattersley, A., Frcp, D., 2016. Prematurity should not prevent genetic testing for neonatal diabetes. Pediatrics 138. 10.1542/peds.2015-3926

19. Estilles, E., Téllez, N., Nacher, M., Montanya, E., 2018. A Model for Human Islet Transplantation to Immunodeficient Streptozotocin-Induced Diabetic Mice. Cell Transplant. 27, 1684–1691. 10.1177/0963689718801006

20. Gibson, D.G., Young, L., Chuang, R.Y., Venter, J.C., Hutchison, C.A., Smith, H.O., 2009. Enzymatic assembly of DNA molecules up to several hundred kilobases. Nat. Methods 2009 65 6, 343–345. 10.1038/nmeth.1318

21. González, B.J., Zhao, H., Niu, J., Williams, D.J., Lee, J., Goulbourne, C.N., Xing, Y., Wang, Y., Oberholzer, J., Blumenkrantz, M.H., Chen, X., LeDuc, C.A., Chung, W.K., Colecraft, H.M., Gromada, J., Shen, Y., Goland, R.S., Leibel, R.L., Egli, D., 2022. Reduced calcium levels and accumulation of abnormal insulin granules in stem cell models of HNF1A deficiency. Commun. Biol. 5. 10.1038/S42003-022-03696-Z

22. Greeley, S.A.W., Polak, M., Njølstad, P.R., Barbetti, F., Williams, R., Castano, L., Raile, K., Chi, D.V., Habeb, A., Hattersley, A.T., Codner, E., 2022. ISPAD Clinical Practice Consensus Guidelines 2022: The diagnosis and management of monogenic diabetes in children and adolescents. Pediatr. Diabetes 23, 1188–1211. 10.1111/PEDI.13426

23. Haataja, L., Arunagiri, A., Hassan, A., Regan, K., Tsai, B., Dhayalan, B., Weiss, M.A., Liu, M., Arvan, P., 2021. Distinct states of proinsulin misfolding in MIDY. Cell. Mol. Life Sci. 78, 6017–6031. 10.1007/S00018-021-03871-1

24. Haataja, L., Manickam, N., Soliman, A., Tsai, B., Liu, M., Arvan, P., 2016. Disulfide mispairing during proinsulin folding in the endoplasmic reticulum. Diabetes 65, 1050–1060. 10.2337/DB15-1345/-/DC1

25. Habeb, A.M., Al-Magamsi, M.S.F., Eid, I.M., Ali, M.I., Hattersley, A.T., Hussain, K., Ellard, S., 2012. Incidence, genetics, and clinical phenotype of permanent neonatal diabetes mellitus in northwest Saudi Arabia. Pediatr. Diabetes 13, 499–505. 10.1111/J.1399-5448.2011.00828.X

26. Herbach, N., Rathkolb, B., Kemter, E., Pichl, L., Klaften, M., Hrabé De Angelis, M., Halban, P.A., Wolf, E., Aigner, B., Wanke, R., 2007. Dominant-Negative Effects of a Novel Mutated Ins2 Allele Causes Early-Onset Diabetes and Severe-Cell Loss in Munich Ins2 C95S Mutant Mice. Diabetes 56, 1268–1276. 10.2337/db06-0658

27. Hrvatin, S., O’Donnell, C.W., Deng, F., Millman, J.R., Pagliuca, F.W., DiIorio, P., Rezania, A., Gifford, D.K., Melton, D.A., 2014. Differentiated human stem cells resemble fetal, not adult, β cells. Proc. Natl. Acad. Sci. U. S. A. 111, 3038–3043. 10.1073/PNAS.1400709111

28. Iafusco, D., Massa, O., Pasquino, B., Colombo, C., Iughetti, L., Bizzarri, C., Mammì, C., Lo Presti, D., Suprani, T., Schiaffini, R., Nichols, C.G., Russo, L., Grasso, V., Meschi, F., Bonfanti, R., Brescianini, S., Barbetti, F., 2012. Minimal incidence of neonatal/infancy onset diabetes in Italy is 1:90,000 live births. Acta Diabetol. 49, 405. 10.1007/S00592-011-0331-8

29. Iafusco, D., Salardi, S., Chiari, G., Toni, S., Rabbone, I., Pesavento, R., Pasquino, B., De Benedictis, A., Maltoni, G., Colombo, C., Russo, L., Massa, O., Sudano, M., Cadario, F., Porta, M., Barbetti, F., 2014. No Sign of Proliferative Retinopathy in 15 Patients With Permanent Neonatal Diabetes With a Median Diabetes Duration of 24 Years. Diabetes Care 37, e181–e182. 10.2337/DC14-0471

30. Iafusco, D., Stazi, M.A., Cotichini, R., Cotellessa, M., Martinucci, M.E., Mazzella, M., Cherubini, V., Barbetti, F., Martinetti, M., Cerutti, F., Prisco, F., 2002. Permanent diabetes mellitus in the first year of life, and the Early Onset Diabetes Study Group of the Italian Society of Paediatric Endocrinology. Diabetologia 45, 798–804. 10.1007/s00125-002-0837-2

31. Ichi Nozaki, J., Kubota, H., Yoshida, H., Naitoh, M., Goji, J., Yoshinaga, T., Mori, K., Koizumi, A., Nagata, K., 2004. The endoplasmic reticulum stress response is stimulated through the continuous activation of transcription factors ATF6 and XBP1 in Ins2 +/Akita pancreatic β β β β cells. Genes to Cells 9, 261–270. 10.1111/j.1365-2443.2004.00721.x

32. Izumi, T., Yokota-Hashimoto, H., Zhao, S., Wang, J., Halban, P.A., Takeuchi, T., 2003. Dominant Negative Pathogenesis by Mutant Proinsulin in the Akita Diabetic Mouse. Diabetes 52, 409–416. 10.2337/DIABETES.52.2.409

33. Johannesson, B., Sui, L., Freytes, D.O., Creusot, R.J., Egli, D., 2015. Toward beta cell replacement for diabetes. EMBO J. 34, 841–855. 10.15252/EMBJ.201490685

34. Kitakaze, K., Oyadomari, M., Zhang, J., Hamada, Y., Takenouchi, Y., Tsuboi, K., Inagaki, M., Tachikawa, M., Fujitani, Y., Okamoto, Y., Oyadomari, S., 2021. ATF4-mediated transcriptional regulation protects against β-cell loss during endoplasmic reticulum stress in a mouse model. Mol. Metab. 54. 10.1016/J.MOLMET.2021.101338

35. Kitamura, Y.I., Kitamura, T., Kruse, J.P., Raum, J.C., Stein, R., Gu, W., Accili, D., 2005. FoxO1 protects against pancreatic beta cell failure through NeuroD and MafA induction. Cell Metab. 2, 153–163. 10.1016/J.CMET.2005.08.004

36. Laybutt, D.R., Preston, A.M., Åkerfeldt, M.C., Kench, J.G., Busch, A.K., Biankin, A. V., Biden, T.J., 2007. Endoplasmic reticulum stress contributes to beta cell apoptosis in type 2 diabetes. Diabetologia 50, 752– 763. 10.1007/S00125-006-0590-Z

37. Lipson, K.L., Fonseca, S.G., Ishigaki, S., Nguyen, L.X., Foss, E., Bortell, R., Rossini, A.A., Urano, F., 2006. Regulation of insulin biosynthesis in pancreatic beta cells by an endoplasmic reticulum-resident protein kinase IRE1. Cell Metab. 4, 245–254. 10.1016/J.CMET.2006.07.007

38. Liu, M., Haataja, L., Wright, J., Wickramasinghe, N.P., Hua, Q.-X., 2010. Mutant INS-Gene Induced Diabetes of Youth: Proinsulin Cysteine Residues Impose Dominant-Negative Inhibition on Wild-Type Proinsulin Transport. PLoS One 5, 13333. 10.1371/journal.pone.0013333

39. Liu, M., Hodish, I., Rhodes, C.J., Arvan, P., 2007. Proinsulin maturation, misfolding, and proteotoxicity. Proc. Natl. Acad. Sci. U. S. A. 104, 15841. 10.1073/PNAS.0702697104

40. Liu, M., Weiss, M.A., Arunagiri, A., Yong, J., Rege, N., Sun, J., Haataja, L., Kaufman, R.J., Arvan, P., 2018. Biosynthesis, structure, and folding of the insulin precursor protein. Diabetes. Obes. Metab. 20, 28. 10.1111/DOM.13378

41. Liu, M., Wright, J., Guo, H., Xiong, Y., Arvan, P., 2014. Proinsulin entry and transit through the endoplasmic reticulum in pancreatic beta cells. Vitam. Horm. 95, 35–62. 10.1016/B978-0-12-800174-5.00002-8

42. Ma, S., Viola, R., Sui, L., Cherubini, V., Barbetti, F., Egli, D., 2018. β Cell Replacement after Gene Editing of a Neonatal Diabetes-Causing Mutation at the Insulin Locus. Stem Cell Reports 11, 1407–1415. 10.1016/J.STEMCR.2018.11.006

43. Maxwell, K.G., Augsornworawat, P., Velazco-Cruz, L., Kim, M.H., Asada, R., Hogrebe, N.J., Morikawa, S., Urano, F., Millman, J.R., 2020. Gene-edited human stem cell-derived β cells from a patient with monogenic diabetes reverse preexisting diabetes in mice. Sci. Transl. Med. 12. 10.1126/SCITRANSLMED.AAX9106

44. Nordmann, T.M., Dror, E., Schulze, F., Traub, S., Berishvili, E., Barbieux, C., Böni-Schnetzler, M., Donath, M.Y., 2017. The Role of Inflammation in β-cell Dedifferentiation. Sci. Reports 2017 71 7, 1–10. 10.1038/s41598-017-06731-w

45. Ortolani, F., Piccinno, E., Grasso, V., Papadia, F., Panzeca, R., Cortese, C., Felappi, B., Tummolo, A., Vendemiale, M., Barbetti, F., 2016. Diabetes associated with dominant insulin gene mutations: outcome of 24-month, sensor-augmented insulin pump treatment. Acta Diabetol. 53, 499. 10.1007/S00592-015-0793-1

46. Oyadomari, S., Koizumi, A., Takeda, K., Gotoh, T., Akira, S., Araki, E., Mori, M., 2002. Targeted disruption of the Chop gene delays endoplasmic reticulum stress–mediated diabetes. J. Clin. Invest. 109, 525. 10.1172/JCI14550

47. Puri, S., Akiyama, H., Hebrok, M., 2013. VHL-mediated disruption of Sox9 activity compromises β-cell identity and results in diabetes mellitus. Genes Dev. 27, 2563. 10.1101/GAD.227785.113

48. Qiao, Z.-S., Guo, Z.-Y., Feng, Y.-M., 2001. Putative Disulfide-Forming Pathway of Porcine Insulin Precursor during Its Refolding in Vitro †. 10.1021/bi001613r

49. Raile, K., O’Connell, M., Galler, A., Werther, G., Kühnen, P., Krude, H., Blankenstein, O., 2011. Diabetes caused by insulin gene (INS) deletion: clinical characteristics of homozygous and heterozygous individuals. Eur. J. Endocrinol. 165, 255–260. 10.1530/EJE-11-0208

50. Redondo, M.J., Hagopian, W.A., Oram, R., Steck, A.K., Vehik, K., Weedon, M., Balasubramanyam, A., Dabelea, D., 2020. The clinical consequences of heterogeneity within and between different diabetes types. Diabetologia 63, 2040. 10.1007/S00125-020-05211-7

51. Renner, S., Braun-Reichhart, C., Blutke, A., Herbach, N., Emrich, D., Streckel, E., Wünsch, A., Kessler, B., Kurome, M., Bähr, A., Klymiuk, N., Krebs, S., Puk, O., Nagashima, H., Graw, J., Blum, H., Wanke, R., Wolf, E., 2013. Permanent Neonatal Diabetes in INS C94Y Transgenic Pigs. Disbetes 62. 10.2337/db12-1065

52. Ron, D., 2002. Proteotoxicity in the endoplasmic reticulum: lessons from the Akita diabetic mouse. J. Clin. Invest. 109, 443–445. 10.1172/JCI200215020

53. Shang, L., Hua, H., Foo, K., Martinez, H., Watanabe, K., Zimmer, M., Kahler, D.J., Freeby, M., Chung, W., LeDuc, C., Goland, R., Leibel, R.L., Egli, D., 2014. β-cell dysfunction due to increased ER stress in a stem cell model of Wolfram syndrome. Diabetes 63, 923–933. 10.2337/DB13-0717

54. Shankar, R.K., Pihoker, C., Dolan, L.M., Standiford, D., Badaru, A., Dabelea, D., Rodriguez, B., Black, M.H., Imperatore, G., Hattersley, A., Ellard, S., Gilliam, L.K., 2013. Permanent Neonatal Diabetes Mellitus: Prevalence and Genetic Diagnosis in the SEARCH for Diabetes in Youth Study. Pediatr Diabetes 14, 174–180. 10.1111/pedi.12003

55. Sharma, R.B., O’Donnell, A.C., Stamateris, R.E., Ha, B., McCloskey, K.M., Reynolds, P.R., Arvan, P., Alonso, L.C., 2015. Insulin demand regulates β cell number via the unfolded protein response. J. Clin. Invest. 125, 3831–3846. 10.1172/JCI79264

56. Snyder, J.T., Darko, C., Sharma, R.B., Alonso, L.C., 2021. Endoplasmic Reticulum Stress Induced Proliferation Remains Intact in Aging Mouse β-Cells. Front. Endocrinol. (Lausanne). 12. 10.3389/FENDO.2021.734079/FULL

57. Son, J., Du, W., Esposito, M., Shariati, K., Ding, H., Kang, Y., Accili, D., 2023. Genetic and pharmacologic inhibition of ALDH1A3 as a treatment of β-cell failure. Nat. Commun. 14. 10.1038/S41467-023-36315-4

58. Stone, S.I., Abreu, D., McGill, J.B., Urano, F., 2021. Monogenic and Syndromic Diabetes due to Endoplasmic Reticulum Stress. J. Diabetes Complications 35, 107618. 10.1016/J.JDIACOMP.2020.107618

59. Støy, J., Edghill, E.L., Flanagan, S.E., Ye, H., Paz, V.P., Pluzhnikov, A., Below, J.E., Hayes, M.G., Cox, N.J., Lipkind, G.M., Lipton, R.B., Atma, S., Greeley, W., Patch, A.-M., Ellard, S., Steiner, D.F., Hattersley, A.T.,Philipson, L.H., Bell, G.I., Performed, A.-M.P., Amemiya, S., Tomita, Y., Darko, D., Doyle, D.A., Densriwiwat, M., Likitmaskul, S., Forsander, G., Hakeem, V., Kocova, M., Liu, L., Macdonald, M.J., Milenkovic, T., Schebek, M., Wentworth, S., 2007. Insulin gene mutations as a cause of permanent neonatal diabetes.

60. Sui, L., Danzl, N., Campbell, S.R., Viola, R., Williams, D., Xing, Y., Wang, Y., Phillips, N., Poffenberger, G., Johannesson, B., Oberholzer, J., Powers, A.C., Leibel, R.L., Chen, X., Sykes, M., Egli, D., 2018a. β-Cell Replacement in Mice Using Human Type 1 Diabetes Nuclear Transfer Embryonic Stem Cells. Diabetes 67, 26–35. 10.2337/DB17-0120

61. Sui, L., Leibel, R.L., Egli, D., 2018b. Pancreatic Beta Cell Differentiation From Human Pluripotent Stem Cells. Curr. Protoc. Hum. Genet. 99. 10.1002/CPHG.68

62. Sui, L., Xin, Y., Du, Q., Georgieva, D., Diedenhofen, G., Haataja, L., Su, Q., Zuccaro, M. V., Kim, J., Fu, J., Xing, Y., He, Y., Baum, D., Goland, R.S., Wang, Y., Oberholzer, J., Barbetti, F., Arvan, P., Kleiner, S., Egli, D., 2021. Reduced replication fork speed promotes pancreatic endocrine differentiation and controls graft size. JCI Insight 6. 10.1172/jci.insight.141553

63. Sun, J., Xiong, Y., Li, X., Haataja, L., Chen, W., Mir, S.A., Lv, L., Madley, R., Larkin, D., Anjum, A., Dhayalan, B., Rege, N., Wickramasinghe, N.P., Weiss, M.A., Itkin-Ansari, P., Kaufman, R.J., Ostrov, D.A., Arvan, P., Liu, M., 2020. Role of proinsulin self-association in mutant INS gene–induced diabetes of youth. Diabetes 69, 954–964. 10.2337/DB19-1106/-/DC1

64. Talchai, C., Xuan, S., Lin, H. V., Sussel, L., Accili, D., 2012. Pancreatic β cell dedifferentiation as a mechanism of diabetic β cell failure. Cell 150, 1223–1234. 10.1016/J.CELL.2012.07.029/ATTACHMENT/8E9EAEEC-C9A5-4769-834E-00CA39266D6B/MMC1.MP4

65. Tanaka, Y., Gleason, C.E., Tran, P.O.T., Harmon, J.S., Robertson, R.P., 1999. Prevention of glucose toxicity in HIT-T15 cells and Zucker diabetic fatty rats by antioxidants. Proc. Natl. Acad. Sci. U. S. A. 96, 10857– 10862. 10.1073/PNAS.96.19.10857

66. Tuch, B.E., Turtle, J.R., Simeonovic, C.J., 1989. Streptozotocin is not toxic to the human fetal B cell. Diabetologia 32, 678–684. 10.1007/BF00274256

67. Wang, J., Takeuchi, T., Tanaka, S., Kubo, S.K., Kayo, T., Lu, D., Takata, K., Koizumi, A., Izumi, T., 1999. A mutation in the insulin 2 gene induces diabetes with severe pancreatic β-cell dysfunction in the Mody mouse. J. Clin. Invest. 103, 27. 10.1172/JCI4431

68. Weiss, M.A., 2009. Proinsulin and the Genetics of Diabetes Mellitus. J. Biol. Chem. 284, 19159. 10.1074/JBC.R109.009936

69. Wright, J., Wang, X., Haataja, L., Kellogg, A.P., Lee, J., Liu, M., Arvan, P., 2013. Dominant protein interactions that influence the pathogenesis of conformational diseases. J. Clin. Invest. 123, 3124–3134. 10.1172/JCI67260

70. Yamada, M., Johannesson, B., Sagi, I., Burnett, L.C., Kort, D.H., Prosser, R.W., Paull, D., Nestor, M.W., Freeby, M., Greenberg, E., Goland, R.S., Leibel, R.L., Solomon, S.L., Benvenisty, N., Sauer, M. V., Egli, D., 2014. Human oocytes reprogram adult somatic nuclei of a type 1 diabetic to diploid pluripotent stem cells. Nature 510, 533–536. 10.1038/nature13287

71. Yong, J., Johnson, J.D., Arvan, P., Han, J., Kaufman, R.J., 2021. Therapeutic opportunities for pancreatic β-cell ER stress in diabetes mellitus. Nat. Rev. Endocrinol. 17, 455. 10.1038/S41574-021-00510-4

72. Yoshinaga, T., Nakatome, K., Nozaki, J.I., Naitoh, M., Hoseki, J., Kubota, H., Nagata, K., Koizumi, A., 2005. Proinsulin lacking the A7-B7 disulfide bond, Ins2Akita, tends to aggregate due to the exposed hydrophobic surface. Biol. Chem. 386, 1077–1085. 10.1515/BC.2005.124/MACHINEREADABLECITATION/RIS

73. Yoshioka, M., Kayo, T., Ikeda, T., Koizumi, A., 1997. A Novel Locus, Mody4, Distal to D7Mitl89 on Chromosome 7 Determines Early-Onset NIDDM in Nonobese C57BL/6 (Akita) Mutant Mice.

74. Yu, M.G., Keenan, H.A., Shah, H.S., Frodsham, S.G., Pober, D., He, Z., Wolfson, E.A., D’Eon, S., Tinsley, L.J., Bonner-Weir, S., Pezzolesi, M.G., King, G.L., 2019. Residual β cell function and monogenic variants in long-duration type 1 diabetes patients. J. Clin. Invest. 129, 3252–3263. 10.1172/JCI127397

