## Supplemental Figure1-8 for "Permanent Neonatal diabetes-causing Insulin mutations have dominant negative effects on beta cell identity"

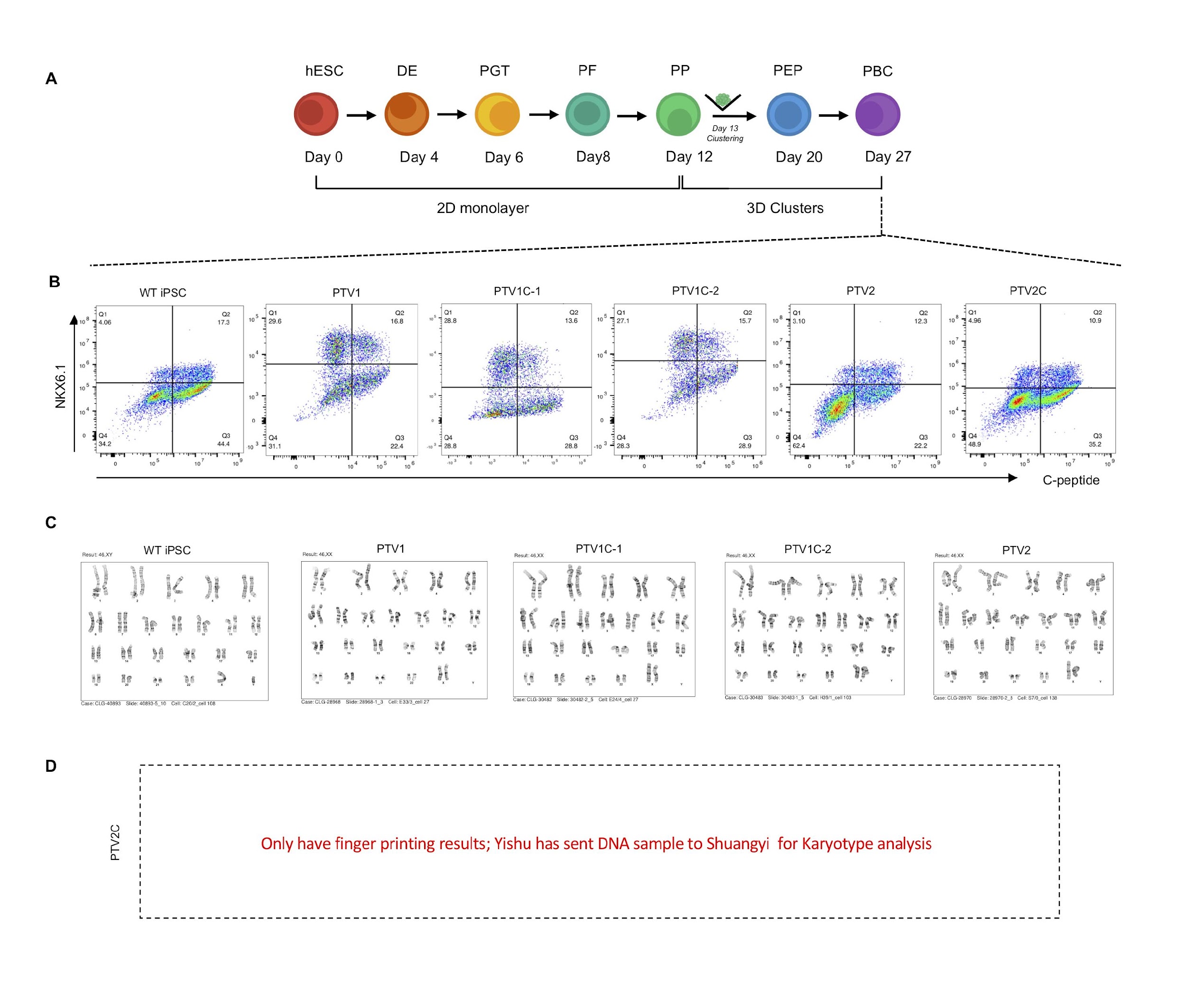

**Figure S1. Related to Figure 1. Characterizations of *in vitro* pancreatic stage SC-beta cells. (A)** Schematic of differentiation protocol mimicking embryonic development of pancreatic β cells *in vitro*. hESC: human embryonic stem cell; DE: definitive endoderm; PGC: primitive gut tube; PF: posterior foregut; PP: pancreatic progenitor; PEP: pancreatic endocrine progenitor; PBC: pancreatic beta cell**.** Until day 13 cells are grown in attachment cultures, dissociated on day 13 and clustered.

**(B)** Pancreatic beta cell clusters (d27) differentiated from WT iPSCs (INS ^+/+^), patient cell lines PTV1(INS ^PTV1/+^), PTV2(INS ^PTV2/+^) and their isogenic controls PTV1C-1(INS ^+/+^), PTV1C-2(INS ^+/+^) and PTV2C(INS ^+/+^). Representative flow cytometry plots of C-peptide and NKX6.1 positive and negative populations of all cell lines. The C-peptide antibody used detects both C-peptide and aa 33-63 of proinsulin.

**(C)** G-band karyotyping of mutant and corrected cell lines.

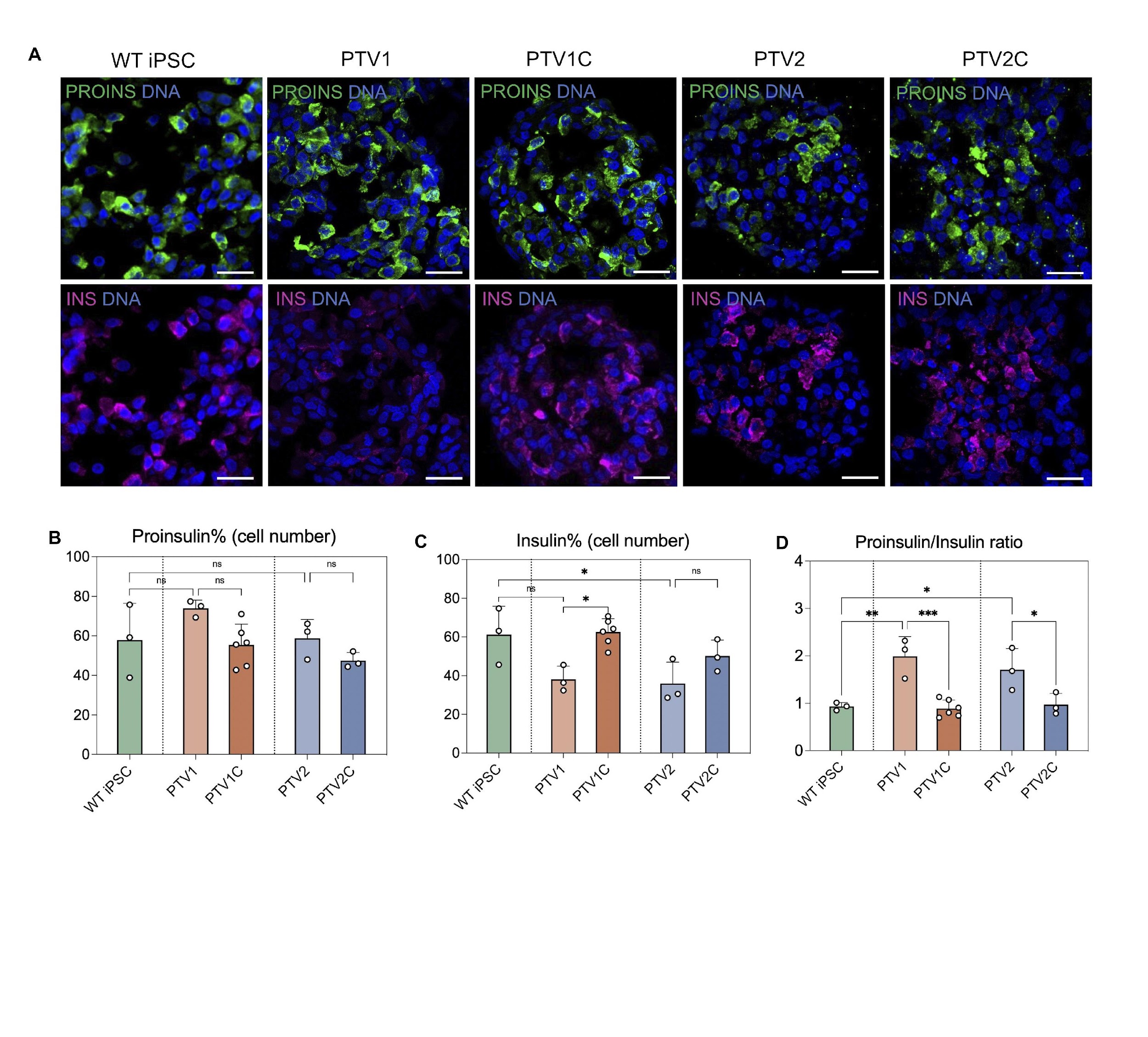

**Figure S2. Related to Figure 2. Reduced proinsulin to insulin process in PTV mutants.**

**(A)** Immunohistochemistry images showing expression of proinsulin, insulin and ER stress marker-BIP in iPSC-derived pancreatic beta cell stage clusters in vitro at d27. scale bar: 20μm.

**(B-C)** Immunofluorescence quantification of proinsulin **(B)** and insulin **(C)** positive cells indicated as percentage of total at a constant exposure time during imaging.

**(D)** Proinsulin-to-insulin ratio (evaluated through the ratio of proinsulin-positive cell number to insulin-positive cell number) representing the proinsulin to insulin processing in iPSC lines. n=3-6 independent experiments per genotype. Data are presented as mean ± SEM. One-way ANOVA or t-test with *P<0.05, **P < 0.01, ***P < 0.001, ****P < 0.0001.

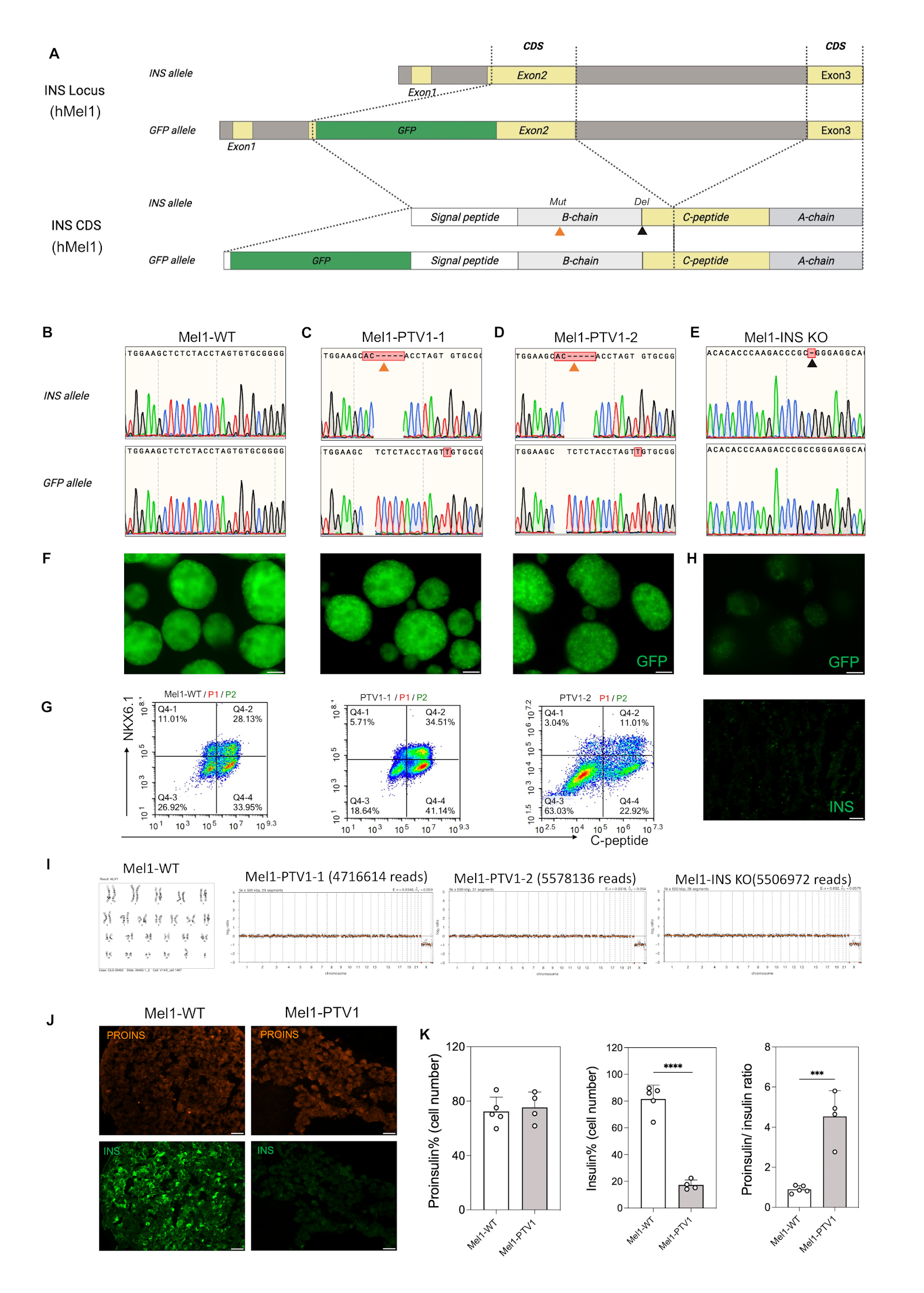

**Figure S3**. **Related to Figure 2. Generation of heterozygous insulin producing cell line with reporter gene.**

**(A)** Schematic and **(B-E)** Sanger sequencing data for Genotypes of WT Mel1 cell line (**Mel1-WT**), Mel1-WT-derived PTV1 mutation cell lines (clone#1: **Mel1-PTV1-1** and clone#2: **Mel1-PTV1-2**) and **(D)** Insulin KO Mel1 cell line (**Mel1-INS KO**).

**(B) Mel1-WT** cell line used in this study is a human embryonic stem cell line with a GFP-coding sequence integrated into the INS locus on one allele (INS ^+/GFP^). The GFP insertion breaks the signal peptide, and therefore, the GFP is intracellular GFP. This reporter line is used in this study for sorting of GFP-positive cells.to facilitate the characterization of insulin-producing (INS+) cells derived *in vitro*.

**(C-D) Mel1-PTV1-1** and **Mel1-PTV1-2** are heterozygous PTV1 mutants generated from Mel1-WT using CRISPR/Cas9, the PTV1 mutation is on the functional insulin allele, and the GFP-inserted allele has an unintentional T insertion at the editing site (Indicated by orange arrows), which does not interfere with GFP production (INS ^PTV1/+^).

**(E) Mel1-INS KO** line is a heterozygous mutant with single C deletion, on the functional insulin allele (Indicated by dark arrow), as the GFP-inserted allele remains unedited.

**(F)** Image of Mel1 and mutant Mel1 cell lines presenting differentiation efficiency indicated by the GFP fluorescence signal.

**(G)** Representative flow cytometry plots for indicated cell lines and markers C-peptide/proinsulin and NKX6.1 on day 27 of differentiation. An antibody recognizing aa33-63 was used. The data are indicative of expression, not processing. The C-peptide antibody used detects both C-peptide and aa 33-63 of proinsulin.

**(H)** Images of Mel1 insulin KO cell line presenting differentiation efficiency indicated by GFP fluorescent signals (Top) and insulin immunofluorescence (Bottom).

**(I)** Karyotype analysis using G-band Karyotyping for Mel1-WT and karyotype analysis using low pass whole genome sequencing for mutant cell lines.

**(J)** Immunohistochemistry images showing expression of proinsulin (Catalog #GNID4) and insulin (Catalog #A0564) in Mel1 derived pancreatic beta cell stage clusters in vitro. Scale bar: 50μm.

**(K)** Immunofluorescence quantitation of proinsulin and insulin are indicated as percentages of all cells detected. Proinsulin-to-insulin ratio (evaluated through the ratio of proinsulin-positive cell number to insulin-positive cell number) representing the proinsulin to insulin processing in Mel1 derived cell lines. n=3-5 frozen sections per genotype. Data plots are presented as mean ± SEM. tt test with *P<0.05, **P < 0.01, ***P < 0.001, ****P < 0.0001.

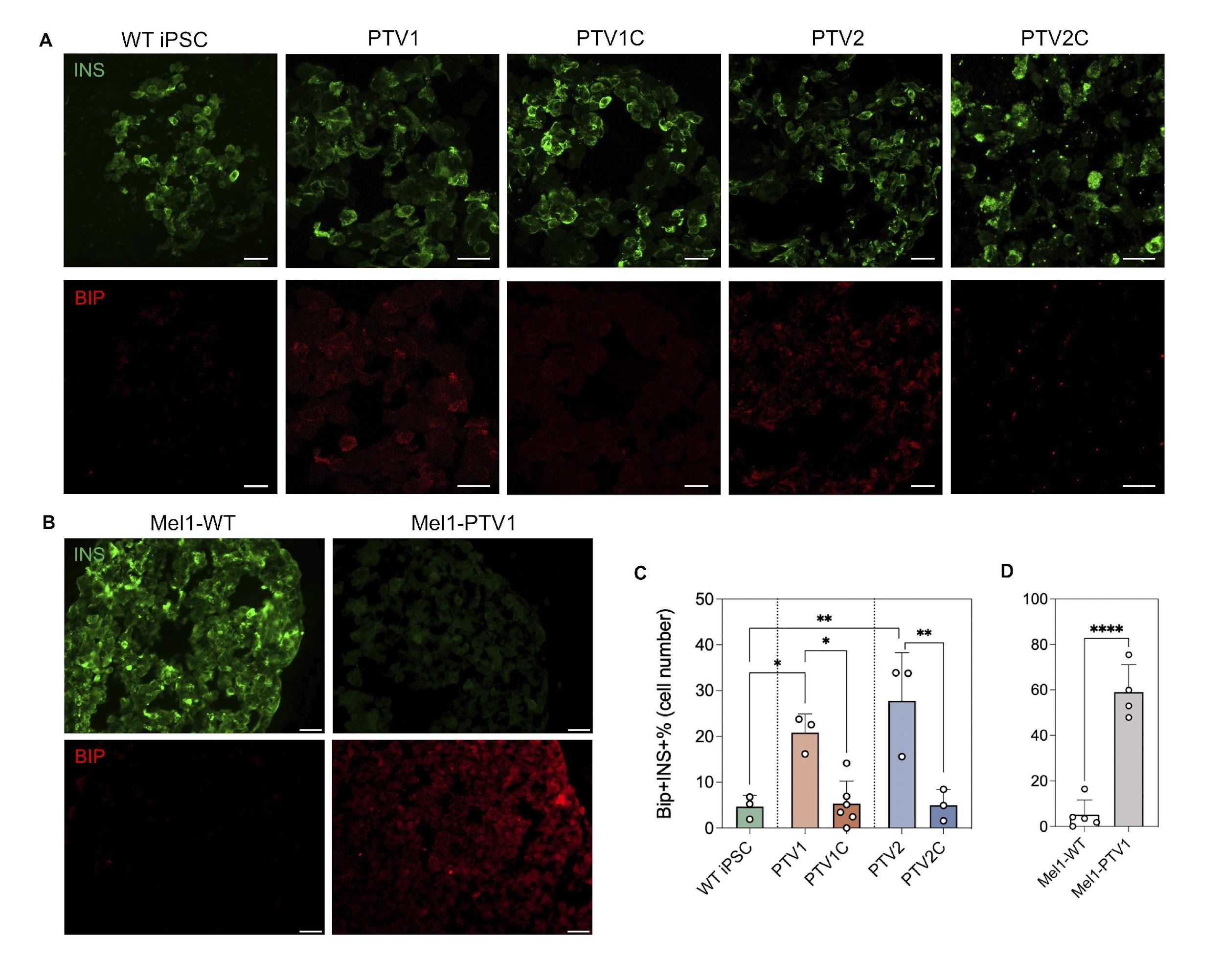

**Figure S4. Related to Figure 2. Elevated ER stress in PTV mutants.**

**(A-B)** Immunohistochemistry images showing expression of Insulin-INS and ER stress marker-BIP in iPSC (A) and Mel1-hESCs (B) derived pancreatic beta cell stage clusters *in vitro*. Scale bar: 20μm in A and 50μm in B.

**(C-D)** Immunofluorescence quantification of INS and BIP double positive percentage in iPSC (C) and Mel1-hESCs (D) derived pancreatic beta cell stage clusters at d27. n=3-5 frozen sections per genotype. Data plots are presented as mean ± SEM. One-way ANOVA test and t test with *P<0.05, **P < 0.01, ***P < 0.001, ****P < 0.0001.

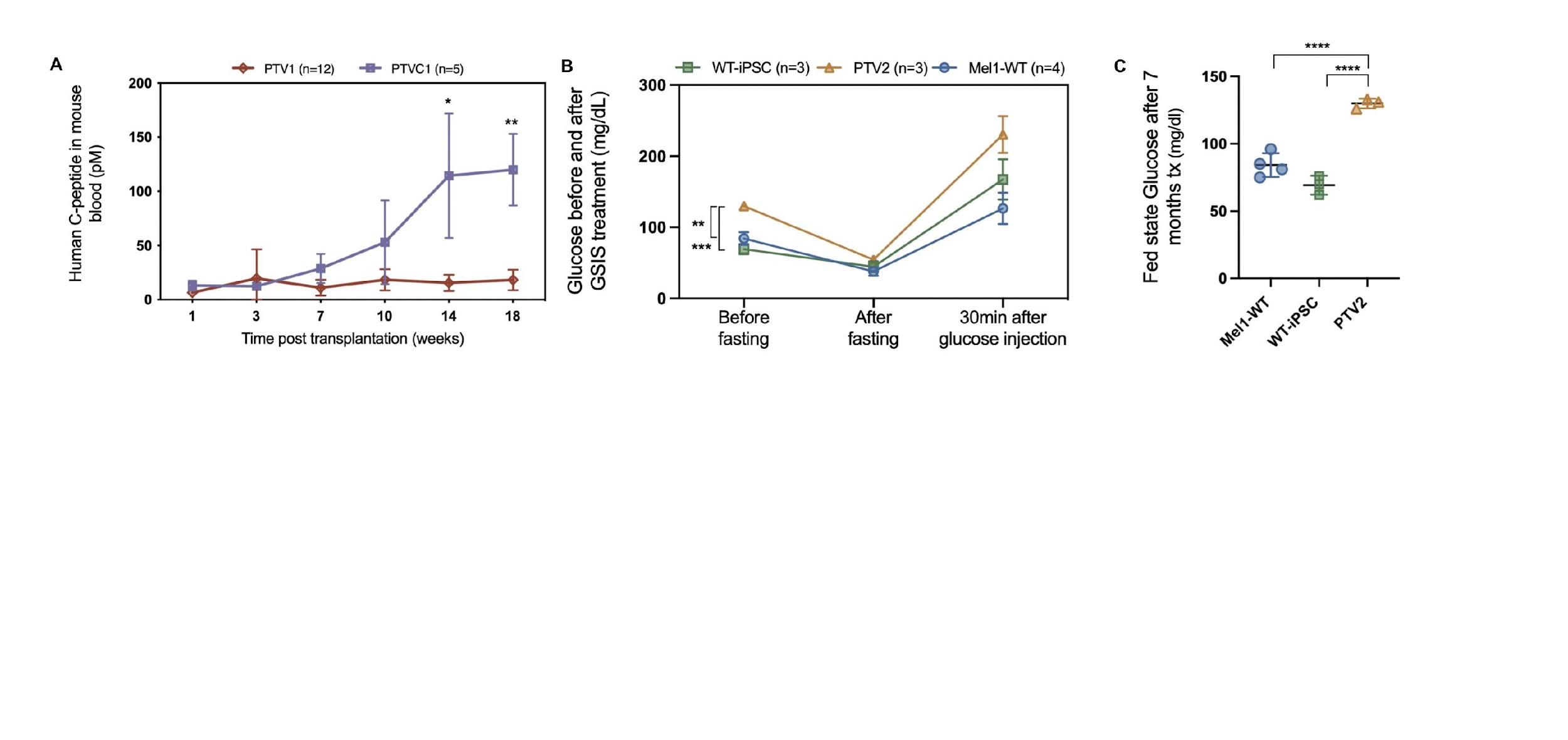

**Figure S5. Related to Figure 3. Transplantation of PTV-edited patient/Mel1-derived SC-beta cells into NGS mice.**

**(A)** Comparisons of human specific C-peptide levels(pM) in PTV1(INS^PTV1/+^) and its isogenic controls PTV1C (INS^+/+^) engraftments. Data plots are presented as mean ± SEM. Two-way ANOVA test with *P<0.05, **P < 0.01, ***P < 0.001, ****P < 0.0001.

**(B)** Blood glucose (mg/dl) levels in mice measured during glucose-stimulated insulin secretion (GSIS) at 7 months post beta cell transplantations. Mice were injected intraperitoneally with 2g of glucose/kg body mass after overnight fasting (16 hours). Data are presented as mean ± SEM. Two-way ANOVA test with *P<0.05, **P < 0.01, ***P < 0.001, ****P < 0.0001.

**(C)** Fed state mouse blood glucose levels(mg/dl) measured at 7 months post-transplantation. Individual data points are shown. One-way ANOVA and Turkey’s multiple comparisons test with *P<0.05, **P < 0.01, ***P < 0.001, ****P < 0.0001.

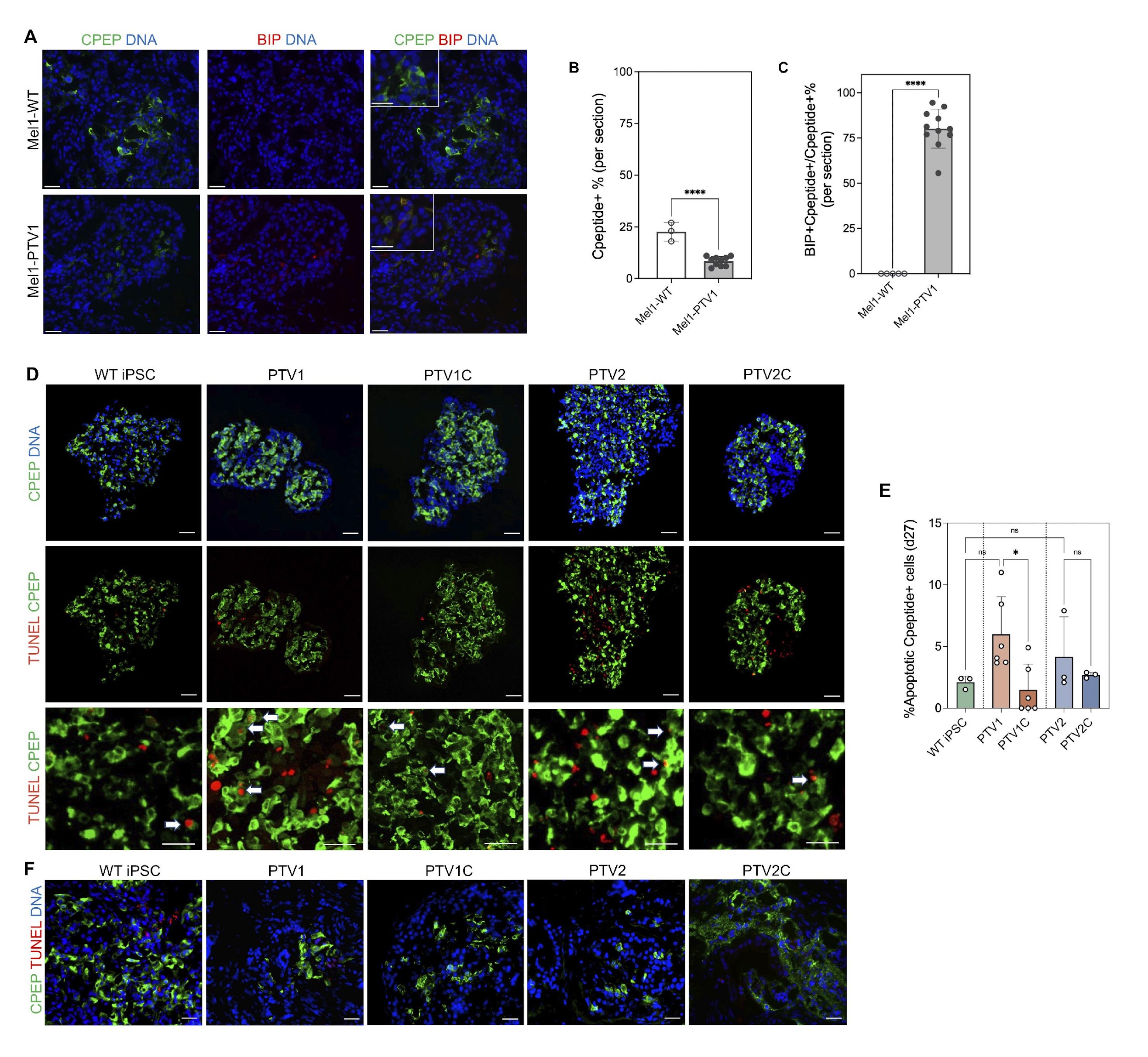

**Figure S6. Related to Figure 4. Early apoptosis in PTV mutants *in vitro* and Increased expression of ER-stress in PTV mutants *in vivo*.**

**(A)** Immunohistochemistry images showing expression of C-peptide (the C-peptide antibody used detects both C-peptide and aa 33-63 of proinsulin) and ER stress marker-BIP in Mel1-WT, Mel1-PTV1 SC-islets in vivo at 7 months post-transplantation. Scale bars = 50um.

**(B)** Quantitation of percentage of C-peptide+ cells.

**(C)** Quantitation of colocalization of C-peptide+ and BIP+. n=2-3 independent transplanted mice per cell genotype. Data were quantified by cell numbers and presented as mean ± SEM. tt test with *P<0.05, **P < 0.01, ***P < 0.001, ****P < 0.0001.

**(D-E)** Apoptotic insulin+ cells assayed by TUNEL staining were increased in PTV1 cells, but not in PTV2 at 27 days in vitro **(D)**, while very rare apoptosis was detected at 7 months post transplantation in vivo **(E)**. Scale bar:50μm.

**(F)** Quantitation of data in (D). n=3-5 frozen sections per genotype. Data plots are presented as mean ± SEM. One-way ANOVA test with *P<0.05, **P < 0.01, ***P < 0.001, ****P < 0.0001.

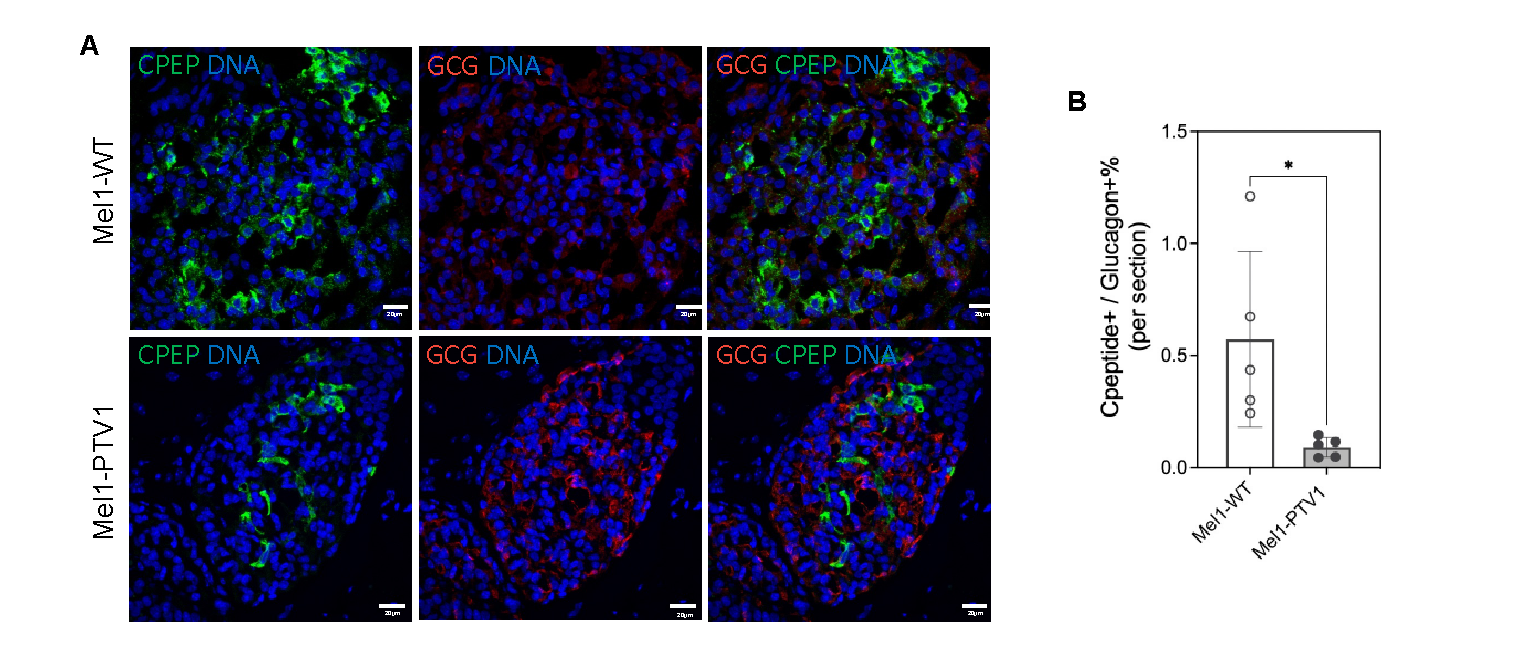

**Figure S7. Related to Figure 5. Reduced C-peptide+/Glucagon+ ratio in Mel1-PTV1 SC-islets engrafted mice.**

**(A)** Immunohistochemistry for beta cells marker C-peptide (the C-peptide antibody used detects both C-peptide and aa 33-63 of proinsulin) and alpha cells marker Glucagon 7 months after transplantation. Scale bar: 20μm.

**(B)** Ratio of insulin producing cell number to Glucagon producing cell number in vivo (7month). n=3 independent transplanted mice per cell genotype. Data were quantified by cell numbers and presented as mean ± SEM. Two-way ANOVA test with *P<0.05, **P < 0.01, ***P < 0.001, ****P < 0.0001.

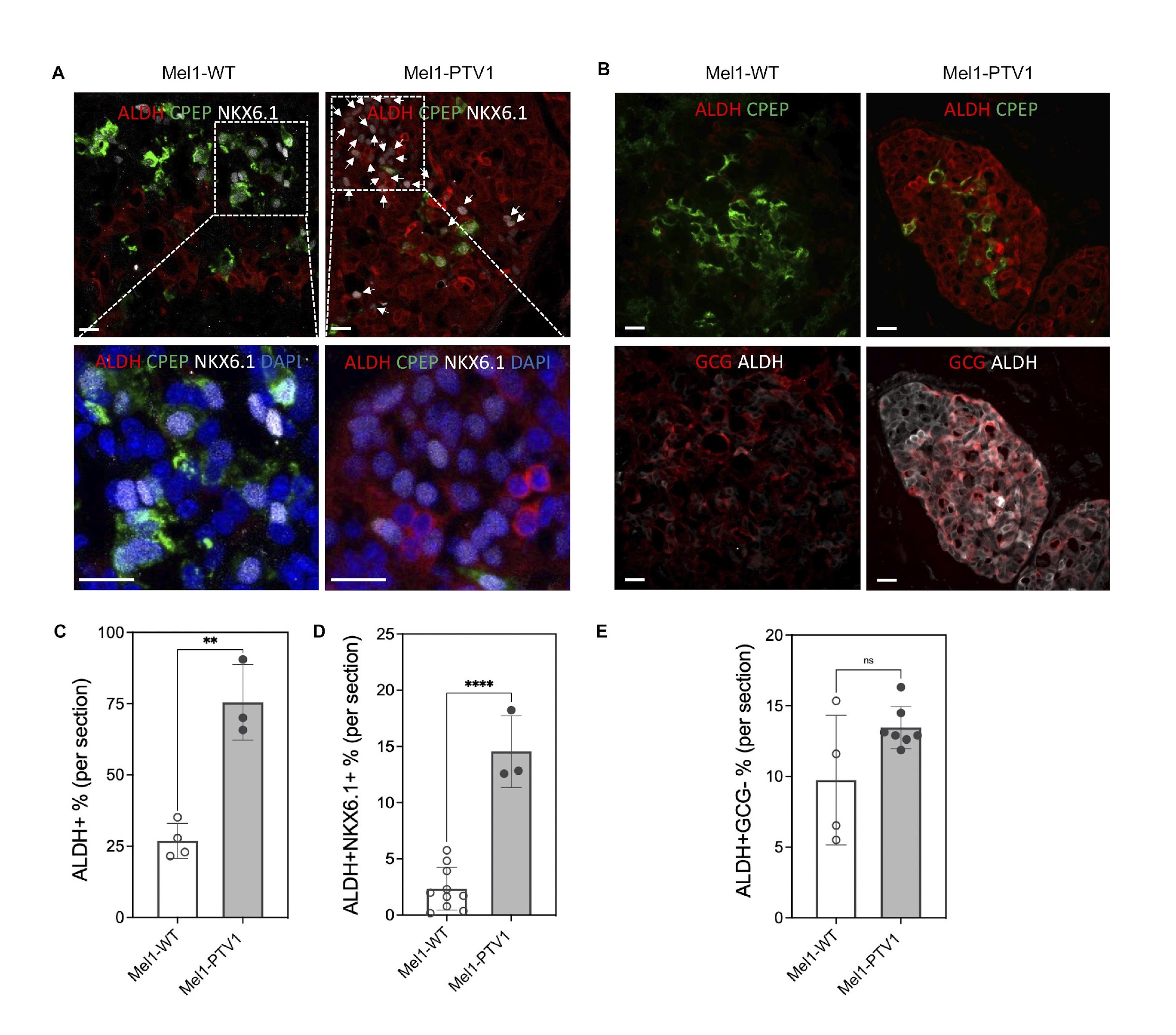

**Figure S8. Related to Figure 6. Increased beta cell dedifferentiation in grafts carrying PTV mutations.**

**(A)** Immunohistochemistry for colocalization of dedifferentiation marker ALDH1A3 (ALDH) with beta cell marker C-peptide (the C-peptide antibody used detects both C-peptide and aa 33-63 of proinsulin), NKX6.1 in 7-month-old grafts. Scale bar: 10μm**.**

**(B)** Percentage of ALDH1A3 positive cells as proportion of all cells.

**(C)** Percentage of NKX6.1 positive cells co-expressing ALDH1A3, indicating dedifferentiated beta cells.

**(D)** Immunohistochemistry for colocalization of dedifferentiation marker ALDH1A3 (ALDH) with C-peptide (the C-peptide antibody used detects both C-peptide and aa 33-63 of proinsulin) and alpha cell marker Glucagon in 7-month-old grafts. Scale bar: 10μm**.**

**(E)** Percentage of GCG negative ALDH1A3 expressing cells. Each data point represents an independent frozen section randomly selected from 3 independent transplanted mice per cell genotype. Data were quantified by cell numbers and presented as mean ± SEM. One-way ANOVA test with *P<0.05, **P < 0.01, ***P < 0.001, ****P < 0.0001.
