## Supplemental Table1-5 for "Permanent Neonatal diabetes-causing Insulin mutations have dominant negative effects on beta cell identity"

**Table S1. Supplemental to Figure1 and Figure 1. Follow up information of PTV2 patient.**

| Time after diabetes onset | Total administered insulin (U/kg/day) | Basal (%) | Bolus (%) | HbA1c (mmol/mol)(%) | C-pep (ng/ml) |
| --- | --- | --- | --- | --- | --- |
| At onset | 0.7 | 88 | 12 | n.r. | 0.47 |
| 6-7 months | 0.57 | 82 | 18 | 50 (6.7) | 0.24 |
| 1 year | 0.69 | 85 | 15 | 43 (6.1) | 0.14 |
| 18 months | 0.67 | 89 | 11 | 44 (6.2) | 0.25 |
| 2 years | 0.7 | 87 | 13 | 39 (5.7) | 0.30 |
| 3 years | 0.83 | 87 | 13 | 50 (6.7) | 0.29 |
| 4 years | 0.9 | 84 | 16 | 46 (6.4) | 0.27 |
| 5 years | 0.84 | 81 | 19 | 44 (6.2) | 0.24 |
| 6 years | 0.81 | 83 | 17 | 48 (6.5) | <0.1 |
| 8 years | 0.82 | 81 | 19 | 57 (7.4) | <0.1 |

**Table S2. Supplemental to Figure1 and Figure 2. Information of human stem cell lines.**

| **Reagent type (species) or resource** | **Designation** | **Source or reference** |
| --- | --- | --- |
| Cell line (Homo sapiens) Male | WT-iPSC | DOI:10.1038/nature13287 |
| Cell line (Homo sapiens) Female | PTV1 | Columbia University Medical Center |
| Cell line (Homo sapiens) Female | PTV1C1 | Columbia University Medical Center |
| Cell line (Homo sapiens) Female | PTV1C2 | Columbia University Medical Center |
| Cell line (Homo sapiens) Female | PTV2 | Columbia University Medical Center |
| Cell line (Homo sapiens) Female | PTV2C | Columbia University Medical Center |
| Cell line (Homo sapiens) Male | Mel1-WT | DOI:10.1007/s00125-011-2379-y |
| Cell line (Homo sapiens) Male | Mel1-PTV1-1 | Columbia University Medical Center |
| Cell line (Homo sapiens) Male | Mel1-PTV1-1 | Columbia University Medical Center |
| Cell line (Homo sapiens) Male | Mel1-INS KO | Columbia University Medical Center |

**Table S3. Supplemental to Figure1 and Figure 2. gRNAs, ssDNA templates and PCR primers used for CRISPR/Cas9 mutations or corrections.**

| **CRISPR/Cas9 Experiments** | **Type** | **Sequence** | **Source or reference** |
| --- | --- | --- | --- |
| PTV1 mutation in WT-Mel1 | gRNA | 5’GAAGCTCTCTACCTAGTGTGCGG3’ | Integrated DNA Technologies |
|  | ssDNA mutation template | 5’CTGACCCAGCCGCAGCCTTTGTGAACCAACACCTGTGCGGCTCACACCTGGTGGAAGCACACCTAGTGTGCGGGGAACGAGGCTTCTTCTACACACCCAAGACCCGCCGGGAGGCAG3’ | Integrated DNA Technologies |
|  | PCR Primers | Forward Primer:  5’ TGCCTCAGCCCTGCCTGTCT3’  Reverse Primer: 5’AAAAGTGCACCTGACCCCCTG3’ | Integrated DNA Technologies |
| *INS* KO in WT-Mel1 | gRNA | 5’GAAGCTCTCTACCTAGTGTGCGG3’ | Integrated DNA Technologies |
|  | PCR Primers | Forward Primer:  5’ TGCCTCAGCCCTGCCTGTCT3’  Reverse Primer: 5’AAAAGTGCACCTGACCCCCTG3’ | Integrated DNA Technologies |
| Correction in PTV1 patient iPSCs | gRNA | 5’GAAGCTCTCTACCTAGTGTGCGG3’ | Integrated DNA Technologies |
|  | ssDNA repair template | 5’CTGACCCAGCCGCAGCCTTTGTGAACCAACACCTGTGCGGCTCACACCTGGTGGAAGCTCTCTACCTAGTGTGCGGGGAACGAGGCTTCTTCTACACACCCAAGACCCGCCGGGAGGCAG3’ | Integrated DNA Technologies |
|  | PCR Primers | Forward Primer:  5’ TGCCTCAGCCCTGCCTGTCT3’  Reverse Primer: 5’AAAAGTGCACCTGACCCCCTG3’ | Integrated DNA Technologies |
| Correction in PTV2 patient iPSCs | gRNA | 5’TCTACACACCCAAGACCCGC3’ | Integrated DNA Technologies |
|  | ssDNA repair template | 5’CTGACCCAGCCGCAGCCTTTGTGAACCAACACCTGTGCGGCTCACACCTGGTGGAAGCTCTCTACCTAGTGTGCGGGGAACGAGGCTTCTTCTACACACCCAAGACCCGCCGGGAGGCAG3’ | Integrated DNA Technologies |
|  | PCR Primers | Forward Primer:  5’ TGCCTCAGCCCTGCCTGTCT3’  Reverse Primer: 5’AAAAGTGCACCTGACCCCCTG3’ | Integrated DNA Technologies |

*Red color represents mutated sequence; Green color represents wildtype sequence.

**Table S4. Primary antibodies used in Flow Cytometry (FC), Immunohistochemistry (IHC) and Western Blot (WB).**

| **Reagent type (species) or resource** | **Designation** | **Identifiers** | **Additional information (used for)** |
| --- | --- | --- | --- |
| Primary antibody | Mouse Anti-NKX6.1 | Catalog #F55A10  Developmental Studies Hybridoma Bank;  RRID: AB_532378 | FC (1:400) IHC (1:100) |
| Primary antibody | Rat Anti-C-peptide/proinsulin (aa 33-63 of proinsulin) | Catalog #GNID4  Developmental Studies Hybridoma Bank;  RRID: [AB_2255626](http://antibodyregistry.org/AB_2255626) | FC (1:100) IHC (1:100). Used for determining differentiation efficiency using flow cytometry. |
| Primary antibody | Mouse Anti-Proinsulin | Catalog #GS9N8  Developmental Studies Hybridoma Bank;  RRID: [AB_532383](http://antibodyregistry.org/AB_532383) | IHC (1:100)  WB (1:500) |
| Primary antibody | Guinea Pig Anti-Insulin | Catalog #A0564  Dako;  RRID: AB_10013624 | IHC (1:1000) |
| Primary antibody | Rabbit Anti-BIP | Catalog #3177S  Cell Signaling Technology;  RRID: AB_2119845 | IHC (1:1000) |
| Primary antibody | Mouse Anti-Glucagon | Catalog #MAB1249-SP  R&D Systems;  RRID: AB_2107340 | FC (1:400) |
| Primary antibody | Rabbit Anti-Glucagon | Catalog # ab92517  Abcam;  RRID: AB_10561971 | IHC (1:1000) |
| Primary antibody | Rabbit Anti-ALDH-A3 | Catalog # NBP2-15339  Novus Biologicals;  RRID: AB_2665496 | IHC (1:1000) |
| Primary antibody | Mouse Anti-KDEL-10C3 | Catalog # ADI-SPA-827  Enzo Life Sciences;  RRID: AB_10618036 | WB (1:1000) |
| Primary antibody | Rabbit Anti-eIF2alpha, phospho (Ser51) | Catalog # 3597  Cell Signaling Technology  RRID: AB_390740 | WB (1:1000) |
| Primary antibody | Mouse Anti-human proinsulin B-C junction sequence KTRREAEDLQ | Abmart  RRID: AB_2921300 | WB (1:1000) |
| Primary antibody | Mouse Anti-Beta Actin | Catalog # CL594-66009  Proteintech;  RRID: AB_2883475 | WB (1:1000) |

**Table S5. Secondary antibodies used Flow Cytometry (FC), Immunohistochemistry (IHC) and Western Blot.**

| **Reagent type (species) or resource** | **Designation** | **Identifiers** | **Additional information** |
| --- | --- | --- | --- |
| Secondary antibody | Donkey anti Rat IgG (H+L), Alexa fluor 488 | Catalog #A-21208  Thermo Scientific;  RRID: AB_141709 | FC (1:500) IHC (1:500) |
| Secondary antibody | Alexa Fluor® 488 Donkey Anti-Rabbit IgG (H+L) Antibody | Catalog #A21206  Life Technologies;  RRID: AB_2535792 | IHC (1:500) |
| Secondary antibody | Alexa Fluor 555 Donkey Anti-Rabbit IgG (H+L) | Catalog #A-31572  Invitrogen;  RRID: AB_162543 | IHC (1:500) |
| Secondary antibody | Alexa Fluor® 555 Donkey Anti-Mouse IgG (H+L) | Catalog #A31570  Life Technologies;  RRID: AB_2536180 | FC (1:500) IHC (1:500) |
| Secondary antibody | Alexa Fluor 647 Donkey Anti-Mouse IgG (H+L) | Catalog #A-31571  Invitrogen;  RRID: AB_162542 | IHC (1:500) |
| Secondary antibody | Alexa Fluor 647 Donkey Anti-Rabbit IgG (H+L) | Catalog # A-31573  Invitrogen;  RRID: AB_2536183 | IHC (1:500) |
| Secondary antibody | Alexa Fluor® 647 AffiniPure Donkey Anti-Guinea Pig IgG (H+L) | Catalog #ab92517  Jackson ImmunoResearch;  RRID: AB_2340476 | IHC (1:500) |
| Secondary antibody | Goat anti-Rat IgG (H+L), Alexa fluor 647 | Catalog #A-21247  Thermo Scientific;  RRID: AB_141778 | IHC (1:500) |
| Secondary antibody | HRP-Goat anti-Mouse | Catalog # 1736516  BioRad  RRID: NA | WB (1:5000) |
| Secondary antibody | HRP-Goat anti-Guinea Pig | Catalog # 1706516, BioRad  BioRad  RRID: NA | WB (1:5000) |
